# Genetic relationships between efficiency traits and gut microbiota traits in growing pigs fed a conventional or a high fiber diet

**DOI:** 10.1101/2021.11.15.468583

**Authors:** V. Déru, A. Bouquet, O. Zemb, B. Blanchet, M.L De Almeida, L. Cauquil, C. Carillier-Jacquin, H. Gilbert

## Abstract

In pigs, the gut microbiota composition plays a major role in the process of digestion, but is influenced by many external factors, especially diet. To be used in breeding applications, genotype by diet interactions on microbiota composition have to be quantified, as well as their impact on genetic covariances with feed efficiency (FE) and digestive efficiency (DE) traits. This study aimed at determining the impact of an alternative diet on variance components of microbiota traits (genera and alpha diversity indices), and estimating genetic correlations between microbiota and efficiency traits for pigs fed a conventional (CO) or a high fiber (HF) diet. Fecal microbes of 812 full-siblings fed a CO diet and 752 pigs fed the HF diet were characterized at 16 weeks of age by sequencing the V3-V4 region of the 16S rRNA gene. A total of 231 genera were identified. Digestibility coefficients of nitrogen, organic matter and energy were predicted analyzing the same fecal samples with near infrared spectrometry. Daily feed intake, feed conversion ratio, residual feed intake and average daily gain (ADG) were also recorded. The 71 genera with less than 20% of zeros were retained for genetic analyses. Heritability (h^2^) of microbiota traits were similar between diets (from null to 0.38 ± 0.12 in the CO diet and to 0.39 ± 0.12 in the HF diet). Only three out of the 24 genera and two alpha diversity indices with significant h^2^ in both diets had genetic correlations across diets significantly different from 0.99 (*P* < 0.05), indicating limited genetic by diet interactions for these traits. When both diets were analyzed jointly, 59 genera had h^2^ significantly different from zero. Based on the genetic correlations between these genera and ADG, FE and DE traits, three groups of genera could be identified. A group of 29 genera was favorably correlated with DE and FE traits, 14 genera were unfavorably correlated with DE traits, and the last group of 16 genera had correlations close to zero with production traits. However, genera favorably correlated with DE and FE traits were unfavorably correlated with ADG, and vice versa. Alpha diversity indices had correlation patterns similar to the first group. In the end, genetic by diet interactions on gut microbiota composition of growing pigs were limited in this study. Based on this study, microbiota-based traits could be used as proxies to improve FE and DE in growing pigs.

## INTRODUCTION

In recent years, gut microbiota has been highlighted as a critical partner of feed and digestive efficiency of pigs, as reviewed by Gardiner et al. (2020) and Patil et al. (2020). Indeed, microbiota composition differs for animals with extreme feed and digestive efficiency values, and some specific intestinal microbes were statistically associated with feed efficiency (McCormack et al., 2017; Yang et al., 2017; McCormack et al., 2019; Bergamaschi et al., 2020b; Aliakbari et al., 2021) and digestive efficiency traits (Niu et al., 2015; Sciellour et al., 2018). Moreover, it has been shown in pigs, as well as in many species, that individual host genetic factors influence the gut microbiota composition. Based on small datasets (< 1000 samples), low to moderate heritability estimates were obtained for certain bacterial genera in pigs (Estelle et al., 2016; Camarinha-Silva et al., 2017; Aliakbari et al., 2021). Recently, a large-scale study in primates with more than 15 000 analyzed samples showed that abundances of most genera are heritable with an average of 6% heritability, indicating that most genera have low heritability (Grieneisen et al., 2021). So far, only one study reported genetic correlations between some heritable genera and feed efficiency traits in growing pigs (Aliakbari et al., 2021). The abundances of some genera were significantly correlated at the genetic level with feed efficiency traits, suggesting opportunities to use microbiota derived traits as proxies of feed efficiency in breeding. Following these results, some authors proposed to use the microbiota information to improve feed efficiency traits (Weishaar et al., 2020; Christensen et al., 2021; Pérez-Enciso et al., 2021).

However, in addition to the genetics of the host, the gut microbiota is affected by many environmental factors, such as diet, and more specifically the amount of dietary fibers in the feed (Zhao et al., 2019; Déru et al., 2021b), and climate (Le Sciellour et al., 2019), internal factors such as sex (Verschuren et al., 2018), as well as uncontrolled microenvironmental effects generally captured by the herd, batch or pen effects (Vigors et al., 2020). Documenting the impacts of these environmental sources of variation on the relationships between microbiota composition and traits is important. Furthermore, a recent GWAS study suggested GxE effects can influence the genetic architecture of microbiota composition traits for pigs reared in a temperate or a tropical environment (Gilbert et al., 2020). However, in this latter study the effect of climate, feed and housing could not be disentangled. To be used in practice, it is critical to estimate the impact of genotype-by-environment (GxE) interactions on the genetic relationships between genera abundances on one hand, and feed and digestive efficiency traits on the other hand.

The objective of this study was to evaluate the genetic relationships between gut microbiota composition and efficiency traits with a focus on the effect of diet on genetic parameters. A large dataset of closely related pigs tested in the same farm with two diets with contrasted dietary fiber contents that significantly affected the abundances of 130 out of 231 genera (Déru et al., 2021b) was used. Genetic parameters of microbiota traits (genera abundances and alpha diversity indices) were estimated within and across diets to quantify the magnitude of genetic by diet interactions. In addition, their genetic correlations with digestive and feed efficiency traits were estimated.

## MATERIAL AND METHODS

The study was conducted in accordance with the French legislation on animal experimentation and ethics. The certificate of Authorization to Experiment on Living Animals was issued by the Ministry of Higher Education, Research and Innovation to conduct this experiment under reference number 2017011010237883 at INRAE UE3P - France Génétique Porc phenotyping station (UE3P, INRAE, 2018. Unité expérimentale Physiologie et Phénotypage des Porcs, France, https://doi.org/10.15454/1.5573932732039927E12).

### Animals and experimental design

Animals considered in this study belonged to the same experimental design as presented in Déru et al. (2020). A total of 1,942 purebred Large White (**LW**) male pigs were reared in 35 successive batches in 2017 and 2018 at the INRAE UE3P France Génétique Porc phenotyping station. A family structure was organized by preferentially testing pairs of full-sibs, each sib being fed one of two dietary sequence that differed in terms of dietary fiber content. Housing conditions and management of pigs were described in Déru et al. (2020). Upon arrival from different nucleus farms, pairs of full-sibs were separated and allotted in pens of 14 animals. Pigs were raised in post-weaning facilities until nine weeks of age and fed with the same standard two-phases post-weaning dietary sequence. Then, they were moved to the growing-finishing facilities without mixing. One group of siblings started to be fed a cereal-based conventional (**CO**) dietary sequence and the other one a high fiber (**HF**) dietary sequence, whose compositions are detailed in the next section. Each growing-finishing pen contained a single place electronic feeder equipped with a weighing scale (Genstar, Skiold Acemo, Pontivy, France) to record feed intake and individual body weight of the animal at each visit to the feeder. At 115 kg body weight, pigs were fasted for 24 hours and then transported to the slaughterhouse. Animals were slaughtered in 89 slaughter batches with an average of 19 pigs per batch. All pigs were issued from 171 sires and 881 dams representative of the French LW collective breeding population.

### Diets

During the growing-finishing phase, the two sets of pigs were fed one of the two-phases dietary sequences. A growing type of diet was first distributed, then a five-days transition was organized at 16 weeks of age and a finishing diet was provided until the end of the test (slaughter body weight of 115kg). The CO diet was formulated to cover energy and amino acids requirements of all pigs. The HF dietary sequence included both soluble dietary fibers and insoluble dietary fibers. The detailed composition of CO and HF feeds is described in Supplementary Table S1. Based on feed formulation, the dietary sequences differed in net energy (**NE**) content, with 9.6 MJ/kg for the CO diet and 8.2 MJ/kg for the HF diet, and in neutral detergent fiber (**NDF**), with 13.90 % for the CO diet and 23.95 % for the HF diet. The ratio digestible Lysine/NE was identical in both dietary sequences, to 0.94 g/MJ NE in the growing phase and to 0.81 g/MJ NE in the finishing phase.

### Measurements and sampling

For each animal, average daily gain (**ADG**), daily feed intake (**DFI**), feed conversion ratio (**FCR**), and residual feed intake (**RFI**) were computed between 35 and 115 kg, namely. The ADG was computed as the ratio between body weight gain and number of days on test. The FCR was calculated as the ratio between DFI and ADG, and was expressed in kg/kg. The RFI was determined using a multiple linear regression of DFI on ADG, lean meat percentage, carcass yield and average metabolic body weight considering data from the two diets jointly in the linear regression, as described in Déru et al. (2020). A spot collection of fecal samples was carried out at 16 weeks of age just before feed transition, for digestive efficiency determination and microbiota composition analyses. For each pig, feces were collected in a piping bag and manually homogenized. To determine digestibility coefficients (**DC**), about 50 g of feces were stored in a plastic container at -20°C until further analyses. Samples were freeze-dried and ground with a grinder (Grindomix GM200, Retsch). Individual DC of energy, nitrogen and organic matter were then predicted based on near infrared spectrometry (**NIRS**) analyses of these samples. Both the methodology to predict DC traits and procedures to validate predictions are described in detail in Déru et al. (2021a). For microbiota DNA extraction, cryotubes (ClearLine cryotubes, Dutscher, France) were filled in with homogenized feces and were immediately frozen in liquid nitrogen. They were then stored at -80°C until analysis.

Pigs that experienced health problems during the test period or had incomplete feed intake data, in equal proportion in the two diets, were discarded from the analysis. In total, 1,663 pigs had FCR, ADG and DFI performances, 1,595 had RFI measurements and 1,242 pigs had NIRS-based predictions of DC traits.

### Microbiota DNA preparation and sequencing

A total of 1,564 fecal samples were used for ribosomal 16S DNA gene sequencing and analysis, with the protocol described in Verschuren et al. (2018). The microbial DNA was extracted using the Quick-DNA Fecal Microbe Miniprep Kit™ (Zymo Research, Freiburg, Germany) and a 15 min bead-beating step at 30 Hz was applied. The V3-V4 region was then amplified from diluted genomic DNA with the primers F343 (CTTTCCCTACACGACGCTCTTCCGATCTTACGGRAGGCAGCAG) and R784 (GGAGTTCAGACGTGTGCTCTTCCGATCTTACCAGGGTATCTAATCCT) using 30 amplification cycles with an annealing temperature of 65 °C. The Flash software v1.2.6 150 (Magoč and Salzberg, 2011) was used to assemble each pair-end sequence, with at least a 10-bp overlap between the forward and reverse sequences, allowing 10% mismatch. Single multiplexing was performed using an in-house 6 bp index, which was added to the R784 primer during a second polymerase chain reaction (**PCR**) with 12 cycles using forward primer (AATGATACGGCGACCACCGAGATCTACACTCTTTCCCTACACGAC) and reverse primer (CAAGCAGAAGACGGCATACGAGAT-index-GTGACTGGAGTTCAGACGTGT). The resulting PCR products were purified and loaded to the Illumina MiSeq cartridge following the manufacturer’s instructions. Run quality was internally checked using PhiX (a library used as a control for Illumina sequencing runs), and each pair-end sequence was assigned to its sample using the bcl2fastq Illumina software. The sequences were submitted to the Short-Read Archive with accession number PRJNA741111. Filtering and trimming of sequences of high quality was applied to the reads with the DADA2 package (Callahan et al., 2016) implemented in R (R Core Team, 2016) using the following parameters: maxN = 0, maxEE = 2, truncQ = 2, trimleft = 17, rm.phix = TRUE and pool = “pseudo”. Chimera were removed with the consensus method to obtain the final operational taxonomic unit (**OTU**) abundance table. No further clustering was applied, so OTUs were equivalent to amplicon sequence variants in this study. This step was followed by taxonomic annotation using the assign Taxonomy function of DADA2 with the Silva Dataset v132 (Quast et al., 2013).

The obtained file was rarefied to 10,000 counts per sample with the Phyloseq R package (McMurdie and Holmes, 2013). A total of 14,366 OTU and 231 genera were finally kept in the abundance tables for 1,564 pigs, including 812 pigs fed the CO diet and 752 pigs fed the HF diet. To facilitate biological interpretations, all analyses were performed at the genus level.

### Statistical analysis

All animals that had at least one data for FCR, ADG, DFI, RFI, DC or microbiota data were kept for following analyses. There were 1,723 animals in total, including 912 animals fed the CO diet and 811 animals fed the HF diet.

### Alpha diversity indices

Based on genera abundances, four alpha diversity indices quantifying the microbial diversity within samples were computed for each animal. First, the richness index was calculated as the number of distinct genera present in the sample. Then, the Shannon index, the Simpson diversity index, and the Chao1 richness estimator were calculated using the following formulas:

1. Chao1 richness estimator: 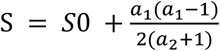
2. Shannon index: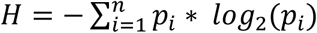
3. Simpson diversity index: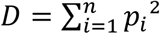 with *p*_*i*_ the abundance of OTU *i, i*= 1,..,*n*, a_1_ the number of OTUs with only one sequence (i.e. ‘singletons’) and a_2_ the number of OTUs with exactly two sequences (i.e. ‘doubletons’) in the sample, and S0 was the observed number of OTUs in the sample. These analyses were performed with the vegan R package (Oksanen et al., 2013).

Genera abundances were normalized using a decimal logarithmic transformation after adding one of each count. For genetic analyses, only genera with less than 20% of zeros in the whole dataset were retained in order to limit deviations from the linear mixed model assumptions, which represented 71 out of the 231 genera. The elementary statistics for these genera are given in Supplementary Table 2 for the whole population and per diet.

### Estimation of variance components within diet

A genetic analysis was undertaken to estimate the host genetic variance for abundances of each genus and each alpha diversity index, as well as their genetic correlations with production traits. A preliminary analysis was carried out to determine fixed and random effects to include in further analyses using a linear mixed model and ignoring additive genetic effects. These analyses were undertaken with the lmer function of the lme4 and lmerTest R packages (Bates et al., 2015; Kuznetsova et al., 2017). Only the effects significant at a threshold of 5% were retained. Then, all traits were analyzed considering each diet separately with the following linear animal mixed model:

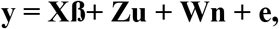

where **y** is the vector of phenotypes for a given trait log10 of genus abundance or alpha diversity index), **ß** is the vector of fixed effects: batch within extraction plate for all genera and alpha diversity indices, and farm of origin for 26 genera. **X** is the incidence matrix relating observations to fixed effects. **Z** is the incidence matrix of the additive genetic effects. **u** ∼ *N*(**0, A** *σ*^*2*^_*u*_) is the vector of additive genetic effects for the considered trait, where **A** is the pedigree relationship matrix built tracing back five generations of pedigree and *σ*^*2*^_*u*_ is the additive genetic variance. **n** ∼ *N*(**0, I***σ*^*2*^_*n*_) is the random effect of the pen nested within batch, applied to all traits. **W** is an incidence matrix relating performances to the random effect **n**. Finally, **e** ∼ *N*(**0, I***σ*^*2*^_*e*_) is the residual random effect. Variance components were estimated by Restricted Maximum Likelihood using the ASREML 4.0 software (Gilmour et al. 2018). Due to potential deviations from normality that may lead to underestimated standard errors, an empirical significance heritability threshold was determined using a bootstrap approach. For two arbitrarily chosen genera, 5,000 univariate analyses were carried out using the aforementioned genetic model after shuffling microbiota abundance values across animals to mimic the null hypothesis of absence of genetic control on genera abundances. Heritability was estimated for each shuffling, and the value of the 95% quantile of the distribution formed by these 5,000 estimations under this null hypothesis was retained as the threshold to decide that the heritability was significantly different from zero, ie that a genus was heritable. Then, to compare heritability estimates across diets, confidence intervals (CI) at 95 and 90% were computed for heritability following two options. First, if the heritability was different from zero (*P* < 0.05), the standard error provided by ASREML was considered as appropriate and the 95% CI was computed as heritability ± 1.96 x standard error, and the 90% CI as heritability ± 1.645 x standard error. Second, when heritability was not different from zero and standard errors could be underestimated, the upper bounds of the 95% CI and 90% CI of a given genera were defined as the sum of the heritability estimate and the corresponding empirical threshold. Heritabilities within non-overlapping CI were considered as significantly different at 90% or 95%, as for a Z-test. In this study, heritability estimates were qualified as low below 0.20, moderate from 0.20 to 0.40, and high above 0.40.

### Estimation of genotype-by-diet interactions

To estimate the magnitude of host genotype by diet interactions on microbiota composition, bivariate analyses were performed across diets for the genera having heritabilities significantly different from 0 in the two diets. To test departure from the null hypothesis of absence of genotype-by-diet interactions (ie genetic correlation between diets close to unity), likelihood ratio tests (LRT) were used to compare a constrained parametric model (H0, genetic correlation = 0.99) with a non-constrained parametric model. The LRT statistic under the null hypothesis then follows the asymptotic distribution of a mixture of a Chi^2^ distribution with 1 degree of freedom and a Dirac (on zero) with equal weights (bordure of the parameter space). The genetic correlations were qualified as low for absolute values between 0.00 and 0.20, moderate between 0.20 and 0.50, and high above 0.50.

### Variance components estimations for both diets combined

To increase statistical power, heritabilities were estimated by combining data from both diets. The same animal mixed model was applied as for separate analyses, adding the fixed effect of the diet, and the random effect of the pen effect within batch was replaced by the pen effect nested within diet nested within batch. The same bootstrapping approach was also used to estimate empirically the 5% significance threshold under the null hypothesis. Then, keeping only microbiota traits with a heritability significantly different from zero, bivariate analyses were carried out to estimate covariances of microbiota traits with ADG and feed and digestive efficiency traits. For the production traits, the animal mixed model included the fixed effects of the diet, of the batch and, as covariates, the weight at the beginning of the test period for ADG, FCR and RFI, the weight at the end of the test period for DFI, and the DFI nested within diet for DC traits to adjust DC for variations due to DFI. Random effects included the effect of the pen nested within batch and diet and the additive genetic effect. The pen within batch and diet effect followed a normal distribution *N*(**0, I***σ*^*2*^_*n*_) and the additive genetic effect followed a normal distribution *N*(**0, A***σ*^*2*^_*g*_) using the same notations and methodology as for microbiota traits. Finally the residual error term followed a normal distribution *N*(**0, I***σ*^*2*^_*e*_) with *σ*^*2*^_*e*_ the residual variance.

A hierarchical cluster analysis was carried out on estimated genetic correlations between gut microbiota traits and production traits, using the Ward distance to identify groups of genera displaying similar patterns of correlations. The clustering was carried out with hclust function of the R package STAT and represented with as.dendogram (Bolar, 2019). The dendogram associated with the heatmap was produced with the heatmap.2 function of the R packages gplots (Warnes et al., 2020). An exact Fisher test was carried out in R (R Core Team, 2016) to identify families overrepresented in the different groups.

Finally, bivariate genetic analyses were carried out to estimate genetic correlations between heritable microbiota traits, using the same linear mixed models and the same methodology as previously presented.

## RESULTS

### Variance components of microbiota traits within diet

Heritability estimated for microbiota traits within each diet are presented along with their 95% confidence interval in Supplementary Table S3. Based on the bootstrap approach, all heritabilities larger than 0.123 in the CO diet and 0.136 in the HF diet were declared significantly different from zero (*P* < 0.05). Heritability of genera abundances were in the same range in the two diets, i.e. from null to 0.38 ± 0.12 in the CO diet and to 0.39 ± 0.12 in the HF diet, respectively. For each genus, heritabilities estimated within each diet are represented in Figure 1. Out of the 71 genera, 24 genera had significant heritability estimates in both diets, 23 genera had significant heritabilities only in one diet, whereas 24 genera were not heritable in any of the diets. None of the 71 genera had heritability estimates significantly different between diets considering 95% confidence intervals. Reducing the confidence intervals to 90% (with 90% empirical thresholds equal 0.09 for the CO diet and 0.10 for the HF diet), heritability estimates of four genera were suggested as different between diets: *Campilobacter, Dialister, Fusicateribacter*, and *Ruminococaceae_UGC002*, the first one having larger h^2^ in the CO diet, whereas the three other ones had larger heritability estimates in the HF diet. Heritability estimates of the four alpha diversity indices were low to moderate and ranged from 0.15 ± 0.09 to 0.23 ± 0.10 in the CO diet, and from zero to 0.20 ± 0.10 in the HF diet. Estimates were not significantly different in the two diets, even if heritability estimates of Chao1 richness estimator and richness were significantly different from zero in the CO diet and not in the HF diet.

**Figure 1.**
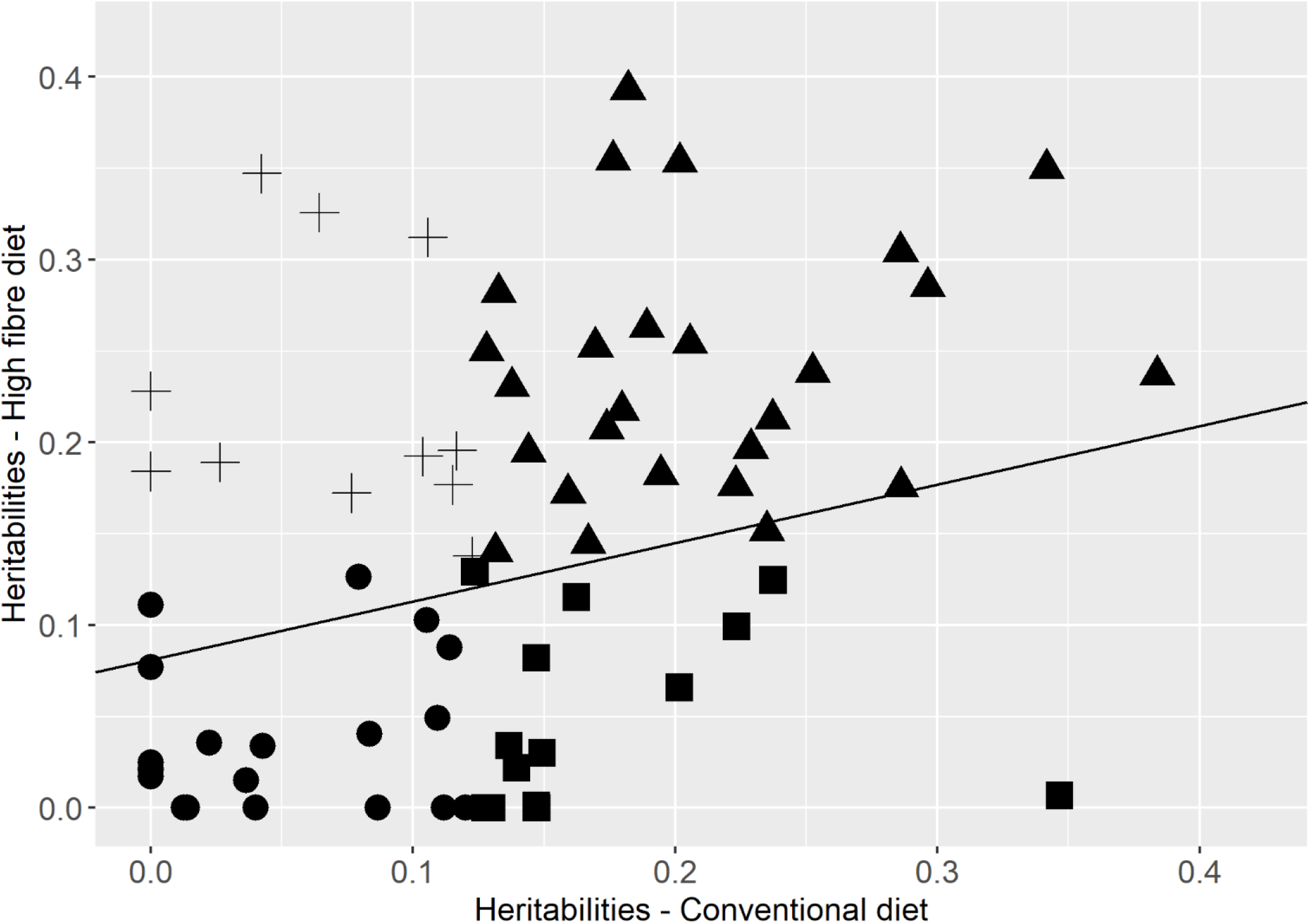
Heritability of the abundances of seventy-one microbial genera, and regression line between estimates determined for growing pigs fed a conventional diet vs pigs fed a high fiber diet. *R*^*2*^ = 0.16. ▴ = genera with heritability significantly different from zero in both diets; • = genera with heritability not significantly different from zero in both diets; ◼ = genera with heritability significantly different from zero in the conventional diet and not in the high fiber diet; + = genera with heritability significantly different from zero in the high fiber diet and not in the conventional diet

### Genetic correlations between diets

Genetic correlations were estimated between the relative abundances of the 24 genera and two alpha diversity indices that had significant heritabilities in both diets (Figure 2 and Supplementary Table S3). Most of these genera had genetic correlations between diets not different from 0.99 according to the LRT, with estimates higher than 0.70 for 18 genera, and between 0.47 ± 0.43 and 0.58 ± 0.38 for three other (*Alloprevotella, Coprococcus_2* and *Ruminiclostridium_6*). Only three genera, *Lachnospiraceae_NK3A20_group* (0.46 ± 0.23), *Mogibacterium* (0.36 ± 0.34) and *Ruminococcus_2* (0.14 ± 0.37), had genetic correlations different from 1. Genetic correlations across diets for the Shannon and Simpson diversity indices, which were both heritable in the two diets, were high and not significantly different from 0.99 (0.83 ± 0.28 and 0.62 ± 0.32).

**Figure 2.**
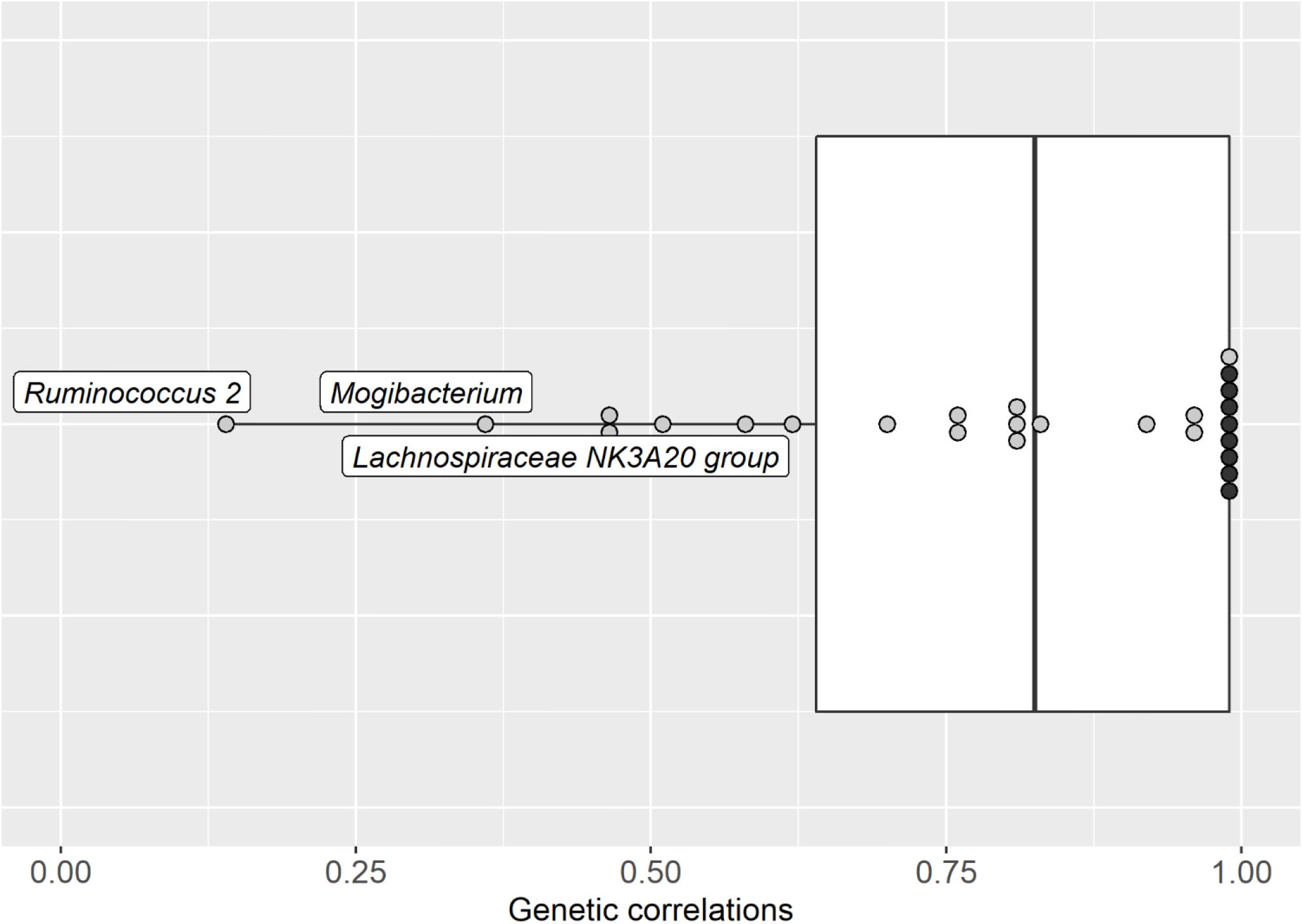
Box-plot of the genetic correlations between diets for microbiota traits (genera and alpha diversity index) with heritability estimates different from zero in each diet, for growing pigs fed a conventional diet and fed a high fiber diet^1,2^ ^1^ The higher the standard errors, the clearer the point. For grey points, standard errors ranged between 0.19 and 0.52. The black points are correlations for which the standard errors were not estimable. ^2^ Name of genera are indicated when genetic correlations significantly differed from 0.99.

### Variance components of microbiota traits for both diets combined

From the previous results, we considered that genetic by diet interactions were limited and microbiota traits could be considered as the same trait in the two diets. Then, heritabilities were estimated for the relative abundances of all genera and diversity indices combining data of both diets to gain statistical power. Resulting heritability estimates, along with their confidence intervals, are presented in Figure 3 and in Supplementary Table S3. Estimated heritabilities were null to moderate (up to 0.31 ± 0.07) and, for most genera, were consistent with those obtained considering the diets separately. Based on the bootstrap approach, all heritabilities higher than 0.058 were significantly different from zero for a 5% type I error. Then, twelve out of the 71 genera had heritability not significantly different from zero, genera that were not heritable in the previous separate analyses. For the 59 heritable genera, 39 genera had low heritability (< 0.20) and 20 genera had moderate heritability (0.20 ≤ h^2^). The most heritable genera were *Anaerovibrio* (0.31 ± 0.07), *Rikenellaceae_RC9_gut* (0.30 ± 0.07), and *Dialister* (0.30 ± 0.06). The three most abundant bacteria in both diets (Supplementary Table S2), *Lactobacillus, Prevotella_9* and *Streptococcus*, had low or moderate heritability (0.18 ± 0.06, 0.12 ± 0.06, and 0.22 ± 0.06, respectively). Estimated heritabilities for the four alpha diversity indices were low (from 0.05 ± 0.04 to 0.19 ± 0.06), and significantly different from zero for all alpha diversity indices but the Chao1 richness estimator.

**Figure 3.**
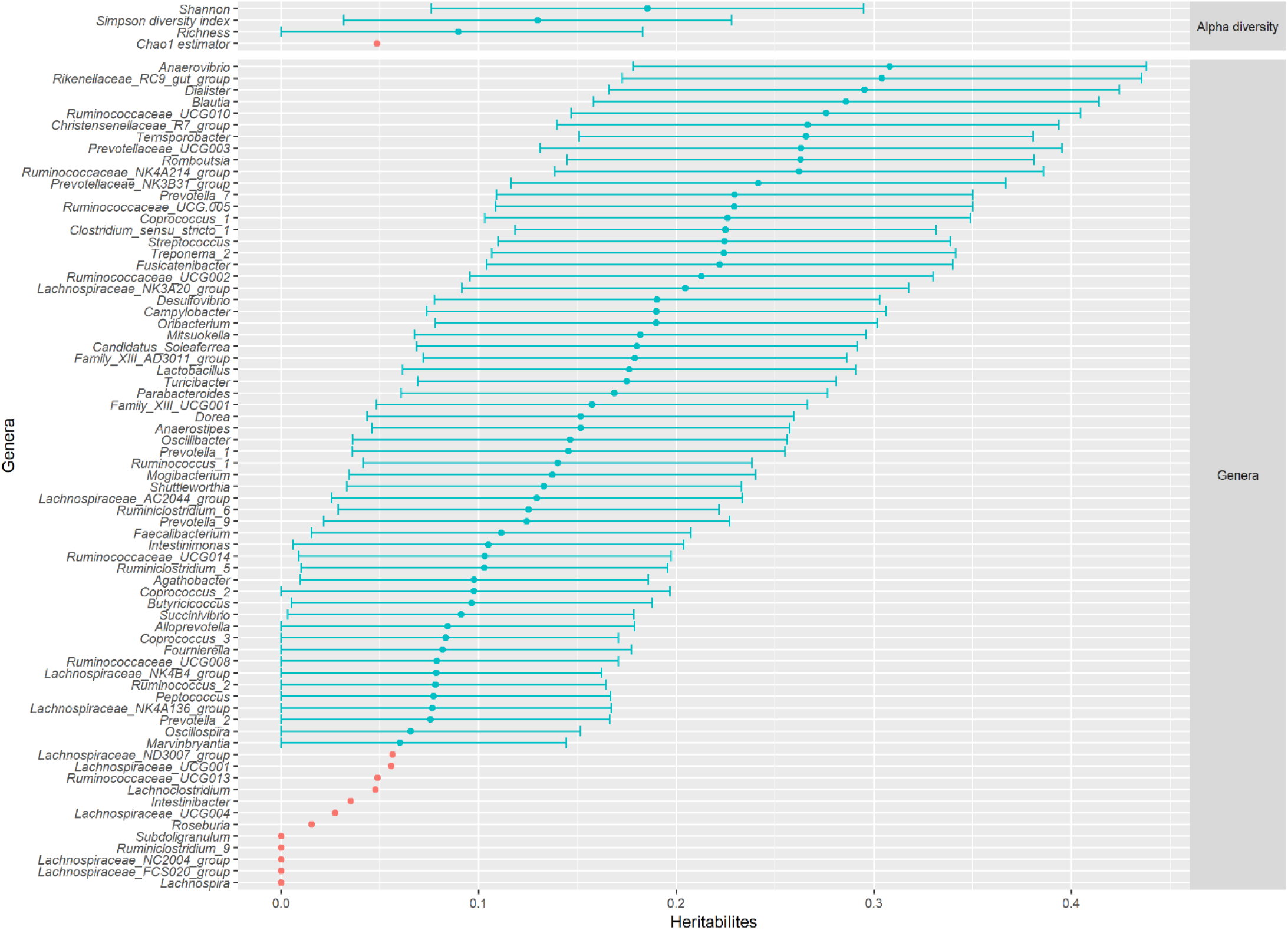
Heritability and 95% confidence intervals of four alpha diversity indices (on top) and seventy-one genera (below) estimated jointly for data of growing pigs fed a conventional diet and growing pigs fed a high fiber diet combined. Orange = heritability not significantly different from zero (*P* > 0.05); blue = heritability significantly different from zero (*P* < 0.05). The 5% threshold was obtained by bootstrapping genera abundances to individuals and determining heritability for each reassignment, with 5000 replicates.

### Genetic correlations with production traits

Genetic correlations between the abundances of the 59 heritable genera on the one hand, and feed and digestive efficiency traits on the other hand, are presented in Figure 4 and Supplementary table S4. After a hierarchical clustering, three groups of genera could be identified depending on their pattern of correlations with production traits (Figure 4). A group of 29 genera was favorably correlated with all traits except ADG, *i*.*e*. had negative correlations with FCR, RFI, DFI, and ADG, and positive correlations with the three DC. A group of 14 genera was unfavorably correlated with digestive and feed efficiency traits, but favorably correlated with ADG, and finally, a group of 16 genera had genetic correlations close to zero with most traits. Based on the Fisher exact test, no family was overrepresented in one of the groups.

**Figure 4.**
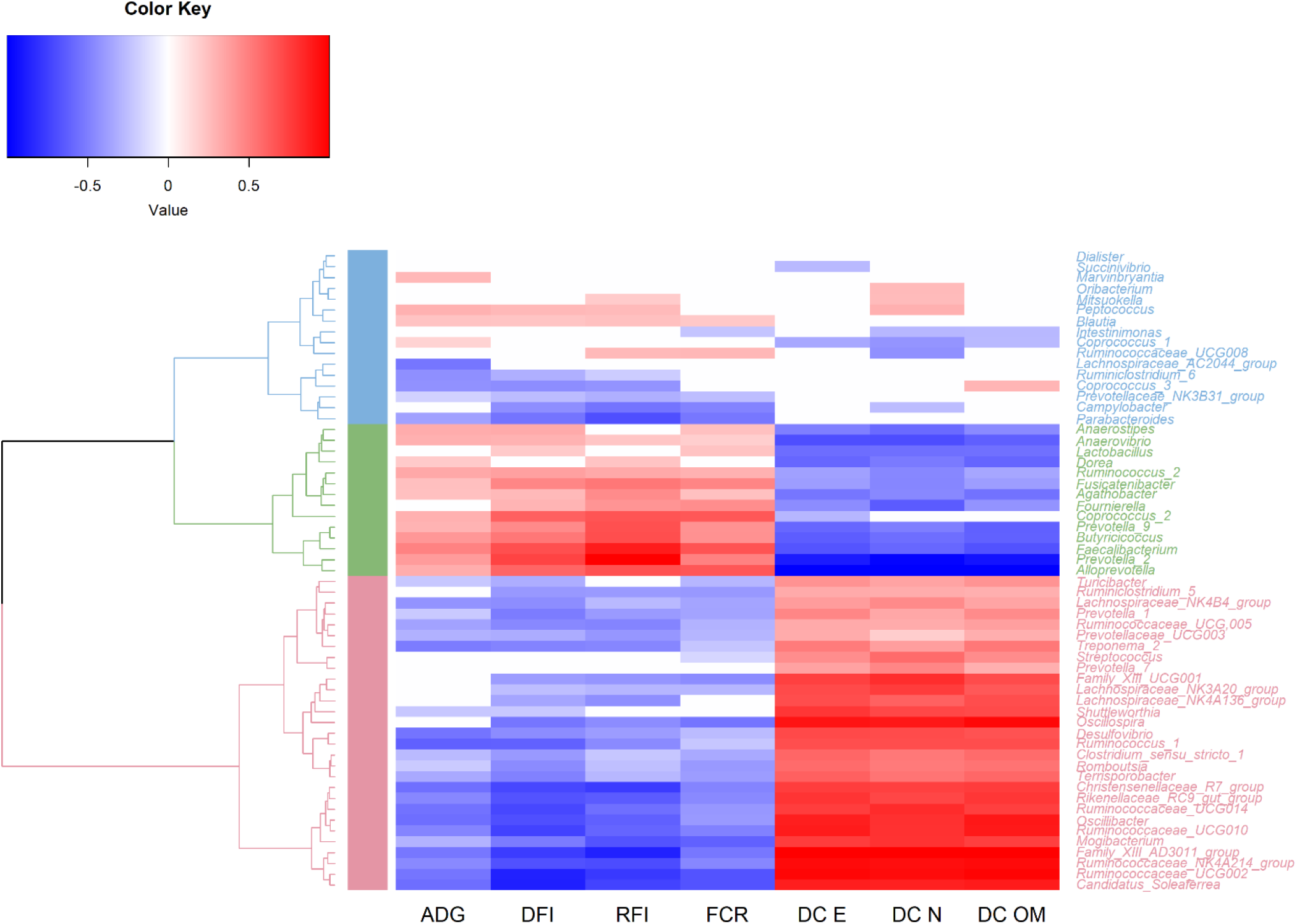
Genetic correlations between abundances of sixty genera with heritability significantly different from zero, and feed and digestive efficiency traits for data of growing pigs fed a conventional and high fiber diet combined^1^. Blue, green and pink colors show limits between groups obtained from the hierarchical analysis of genetic correlations between these genera and production traits. ADG = average daily gain; DFI = daily feed intake; RFI = residual feed intake; FCR= feed conversion ratio; DC E = digestibility coefficient of energy; DC N = digestibility coefficient of nitrogen; DC OM = digestibility coefficient of organic matter ^1^The 68% confidence interval was determined by subtracting or adding one standard error to the estimated heritability. If zero was included within this confidence interval then heritability was set to zero

Genetic correlations between the three alpha diversity indices, and feed and digestive efficiency traits are presented in Table 1. In general, these indices were negatively correlated with ADG, DFI, RFI and FCR, *i*.*e*. were favorably correlated with all traits except ADG, and positively, *i*.*e*. favorably, correlated with DC traits. This pattern corresponded to the one observed for the first group of genera described above. However, there were differences of correlation magnitudes between the Shannon and Simpson indices on the one hand, and the richness on the other hand. The Shannon and Simpson diversity indices had low correlations, not significantly different from zero, with ADG, moderate to high correlations with DFI, RFI and FCR (from -0.56 ± 0.13 to -0.42 ± 0.17), and very high correlations with DC traits (> 0.82 ± 0.12). The richness was strongly and significantly correlated with ADG and DFI, and moderately to highly correlated with the other traits.

**Table 1.** Genetic correlations between three alpha diversity indices with heritability significantly different from zero, and feed and digestive efficiency traits for data of growing pigs fed a conventional and high fiber diet combined, along with their standard errors

|  | Feed efficiency traits |  |  |  | Digestive efficiency traits |  |  |
| --- | --- | --- | --- | --- | --- | --- | --- |
|  | ADG | DFI | RFI | FCR | DC E | DC N | DC OM |
| Shannon index | -0.29<br>± 0.17 | -0.56<br>± 0.13 | -0.51<br>± 0.13 | -0.46<br>± 0.14 | 0.91<br>± 0.13 | 0.88<br>± 0.12 | 0.90<br>± 0.13 |
| Simpson index | -0.19<sup>1</sup><br>± 0.19 | -0.48<br>± 0.16 | -0.42<br>± 0.17 | -0.42<br>± 0.17 | 0.99<br>± NE<sup>2</sup> | 0.99<br>± NE<sup>2</sup> | 0.99<br>± NE<sup>2</sup> |
| Richness | -0.87<br>± 0.27 | -0.81<br>± 0.20 | -0.61<br>± 0.20 | -0.55<br>± 0.18 | 0.74<br>± 0.24 | 0.58<br>± 0.24 | 0.70<br>± 0.24 |

ADG = average daily gain; DFI = daily feed intake; RFI = residual feed intake; FCR= feed conversion ratio; DC E = digestibility coefficient of energy; DC N = digestibility coefficient of nitrogen; DC OM = digestibility coefficient of organic matter
<sup>1</sup>Correlation not significantly different from zero according to the 68% confidence interval determined by subtracting and adding one standard error to the estimated heritability.
<sup>2</sup> NE= Not estimable

### Genetic correlations between genera and diversity indices

Genetic correlations estimated between genera are represented in Figure 5, keeping the genera ranking from the hierarchical clustering as in Figure 4. Most genera belonging to the first group, the 29 genera favorably correlated with digestive and feed efficiency traits, were positively correlated with each other. Similarly, the 14 genera within the group of genera favorably correlated with growth rate but unfavorably with feed and digestive efficiency were generally positively correlated with each other. The genetic correlations between these two groups were mostly negative. Finally, no clear correlation pattern could be identified between genera belonging to the group of genera weakly correlated with the production traits.

**Figure 5.**
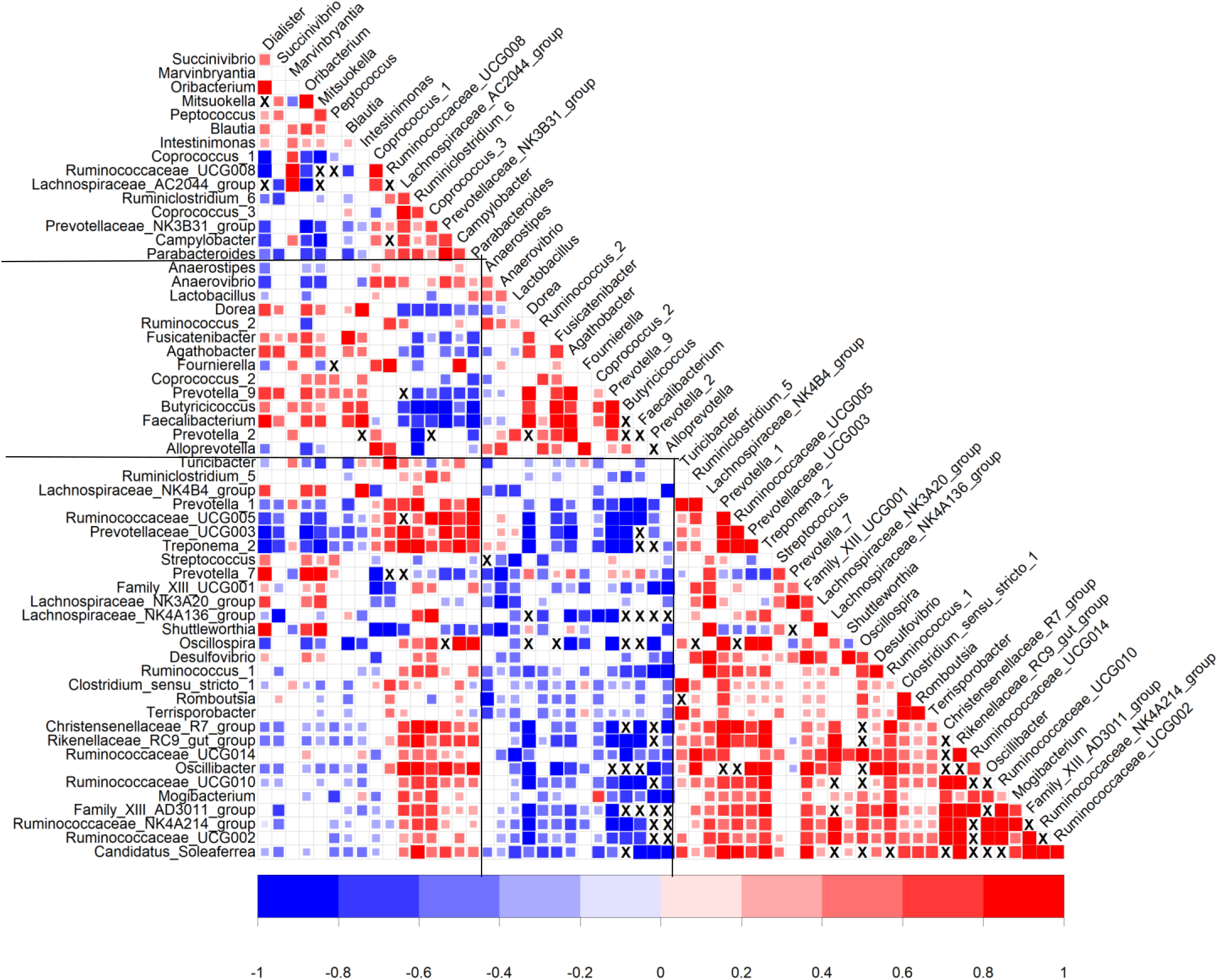
Genetic correlations^1^ between sixty heritable genera considering data of growing pigs fed the conventional or high fiber diets. Vertical bars show limits between groups obtained from the hierarchical analysis of genetic correlations between these genera and production traits. ^1^Correlations were represented in colors when they were significantly different from zero considering alpha=32%, and were left blank otherwise. Genetic correlations that could not be estimated due to convergence problems of the algorithm are marked with an X.

Finally, the genetic correlation between Shannon and Simpson diversity indices was estimated to 0.95 ± 0.03. Genetic correlations estimated between the Shannon and Simpson diversity indices on one hand and genera abundances on the other hand are presented in Figure 6. Most genera from the groups associated with higher digestive efficiency were moderately to strongly correlated with diversity indices. On the contrary, the genera associated with higher growth rate and poorer digestive efficiency were unfavourably genetically correlated with gut microbiota diversity indices. Genera that were weakly correlated both with growth rate and digestive efficiency were also weakly to moderately correlated with diversity indices.

**Figure 6.**
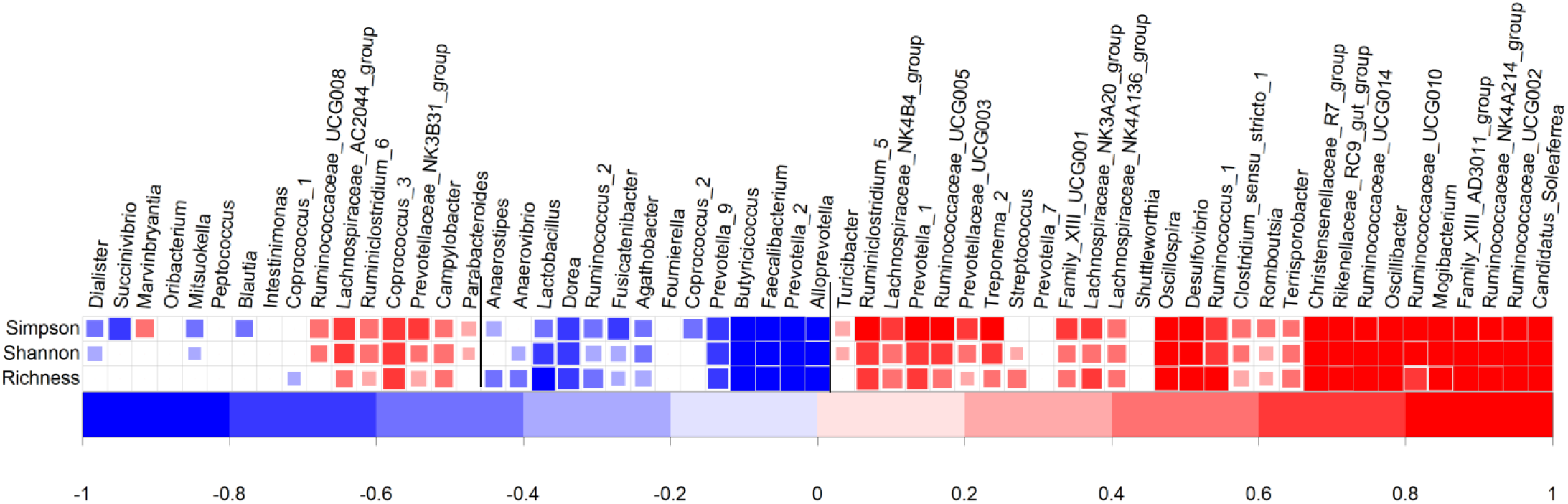
Genetic correlations^1^ between gut microbiota diversity indices and genera abundances considering data of growing pigs fed the conventional or high fiber diets. Vertical bars show limits between groups obtained from the hierarchical analysis of genetic correlations between these genera and production traits. ^1^Correlations were represented in colors when they were significantly different from zero considering alpha=32%, and were left blank otherwise.

## DISCUSSION

From a phenotypic point of view, it is acknowledged that the diet has an impact on the microbiota composition in growing pigs (Bauer et al., 2006; Verschuren et al., 2018), which was also observed in the present experimental design (Déru et al., 2021b). Results presented in this study confirmed that microbial abundances and microbial diversity measures are lowly to moderately heritable, as previously shown in pigs (Estelle et al., 2016; Camarinha-Silva et al., 2017; Lu et al., 2018; Bergamaschi et al., 2020a; Aliakbari et al., 2021) and other livestock species (Difford et al., 2018). Based on the subset of genera that were significantly heritable in both diets, our findings suggest limited genotype by diet interactions for most genera. Finally, some microbiota traits were genetically correlated with digestive and feed efficiency traits, making them interesting in a breeding perspective.

### Impact of the diet on variance components estimates of microbiota traits

Impact of the diet on variance components of the microbiota traits in this study could have been underestimated due to the limited size of the datasets within diet (812 and 752). However, heritability of microbiota traits were low to moderate within diet, and similar to those reported in the literature (Estelle et al., 2016; Camarinha-Silva et al., 2017; Aliakbari et al., 2021). Heritability estimated for each genus and alpha diversity indices were generally similar between diets. Thus, even though mean abundances were significantly different for 66 out of the 71 genera in this dataset (Supplementary table S2 Déru et al., 2021b), the proportion of phenotypic variance under genetic control was similar in both diets. This also holds for alpha diversity indices, with higher diversity indicators in the HF diet (3.14 ± 0.23 vs 3.06 ± 0.26 for the Shannon index for instance, *P* < 0,001) but similar heritability estimates.

In our study, genetic correlations estimated between microbiota traits across diets were generally high (≥ 0.70), suggesting that genotype by diet interactions were limited for most of them. Out of the 24 considered genera, only *Lachnospiraceae_NK3A20_group, Mogibacterium* and *Ruminococcus_2* had correlations different from 1. Their heritability estimates were significantly different from zero, and similar in the two diets. In our study, pigs fed a HF diet had a lower abundance of the *Mogibacterium* genus than pigs fed the CO diet (*P* < 0.001). Zhu et al. (2020) showed a positive correlation between the abundance of the *Mogibacterium* genus and the concentration of short-chain fatty acid (SCFAs) in feces of pigs at 42 days of age. SCFAs are molecules (such as butyrate, acetate and proprionate) produced by bacteria when they digest fibers (Slavin, 2013). Accordingly, pigs fed a high fiber diet are expected to have a significantly higher abundance of the *Mogibacterium* genus, which this was not the case in our results. However, in our study feces were collected later than Zhu et al. (2020) (112 days vs 42 days), which could explain the lower abundance of *Mogibacterium* genus at this physiological stage. The abundances of *Ruminococcus_2* and *Lachnospiraceae NK3A20* genera were significantly higher for pigs fed the CO diet than pigs fed the HF diet (*P* < 0.03). Ze et al. (2012) showed that *Ruminococcus* was involved in the degradability in resistant starch in the human colon. Helm et al. (2021) observed for sows an increase of the abundance of *Lachnospiraceae NK3A20* when an intermediate fermentability diet (comparable to the CO diet of our experiment) was given to the pigs compared to low fermentable diet and highly fermentable diet. All diets in the experiment of Helm et al. (2021) were inoculated with *Brachyspira hyodysenteriae*, a bacteria causing diarrhea. Nevertheless, these comparisons with the literature are made difficult by taxonomy nomenclature differences (in recent databases *Ruminococcus 1* and *Ruminococcus 2* are distinguished), and different experimental conditions (diet, date of feces collection, limited number of data…). Further studies to better understand the metabolic interplay of these microbiota and the host in different dietary conditions would be necessary. However, altogether a diet with increased fiber content seems to have a limited impact on the variance components of the microbiota traits. Consistent conclusions have been drawn for growth rate, feed and digestive efficiency traits based on this experiment: feeding pigs with the high fiber diet significantly affects mean performances, but would imply limited genotype by diet interactions that would not affect breeding strategies (Déru et al. 2020; Déru et al. 2021a).

### Genetic Associations Between Gut Microbiota and Digestive Efficiency Traits

#### Clustering Genera

In our study, three groups of genera were highlighted, with very consistent relationships with production traits: those favorably correlated, those unfavorably correlated, and those with very low or null correlations. These correlations were consistent with the genetic correlations between production traits and alpha diversity indexes. This high consistency among microbiota traits is certainly due to the high structuration of microbiota populations, as shown by the genetic correlation matrix among the microbiota traits: microbiota within each of the two groups showed high positive correlations to each other, and negative correlations with genera from the other group. Rather than analyzing all genera independently, enterotypes could be identified, as previously done in growing pigs. In a longitudinal study, Ramayo-Caldas et al. (2016) identified two main enterotypes five weeks after weaning. Those enterotypes were dominated either by *Ruminococcus* and *Treponema*, or by *Prevotella* and *Mitsuokella*. Such definition of enterotypes could be consistent with the present results. The first group of genera associated with higher digestive efficiency comprises many genera of the *Ruminococcaceae* family as well as *Treponema_2* and was strongly correlated with alpha diversity indices, in agreement with Ramayo-Caldas et al. (2016), but also with Le Sciellour et al. (2019) who defined other enterotypes in older pigs submitted to heat stress. However, the studies of Ramayo-Caldas et al. (2016) and Le Sciellour et al. (2019) reported different enterotypes at different ages and environmental conditions, Bergamaschi et al. (2020b) reported different enterotypes in different breeds, and those identified by Le Sciellour et al. (2019) did not show phenotypic correlations with production traits, so these approaches might also show limits for breeding.

#### Validation with other studies

So far, only few studies have reported genetic relationships and correlations between microbiota and production traits. In the recently published study of Aliakbari et al. (2021), microbiota differences between pig lines divergently selected for RFI were computed. Their dataset was independent from ours (different animals, farms and diets), but the same molecular procedures and bioinformatic pipelines were used to produce the microbiota data. Among the six genera most favorably associated with digestive and feed efficiency in our study, five were more abundant in their low RFI line (*Candidatus_Soleaferrea* was too rare to be analyzed), whereas among the six genera most unfavorably associated with digestive and feed efficiency in our study, five were less abundant in their low RFI line (*Alloprevotella, Prevotella_2, Faecalibacterium, Prevotella_9* and *Coprococcus_2*), and one showed no line difference (*Butyricicoccus*). In addition, in Aliakbari et al. (2021), three genera were significantly correlated with RFI at the genetic level, including *Streptococcus* that was not correlated with efficiency traits in our study and was unfavorable in Aliakbari et al. (2021). For the two other genera, *Prevotella 2* and *Desulfovibrio*, our results showed the same directions of correlations with RFI as in Aliakbari et al. (2021): *Prevotella 2* belonged to the second group based on correlation, ie was unfavorably correlated with feed efficiency, and *Desulfovibrio* to the first group. A similar consistency was observed for the genera found genetically associated with DFI in Aliakbari et al. (2021): four of them showed the same directions of correlations in our study, and the three others were not heritable. Furthermore, the three alpha diversity indices with significant heritabilities were favorably correlated with digestive efficiency traits and, as previously reported by Aliakbari et al. (2021), with feed efficiency traits. It has been hypothesized that increase in the number of different microbes may improve absorption of nutrients via providing flexibility and redundancy to the microbiota to exploit various sources of nutrients (Moya and Ferrer, 2016) and contributing to gut health (Fouhse et al., 2016). Our results were thus very consistent with their report. Compared with the recent review from Gardiner et al. (2020) which pointed out many discrepancies between studies reporting microbiota-production relationships, our approach suggests that standardization of the procedures to build microbiota data, including use of amplicon variant sequences instead of clustered OTU as recommended by Callahan et al. (2017), would greatly facilitate comparisons across microbiota studies.

#### Relationships between efficiency, growth rate and microbiota traits

With growth rate, the signs of the correlations were similar to that of efficiency traits, with reduced magnitudes of the correlation estimates, leading to antagonistic correlations of microbiota traits with efficiency and growth. Aliakbari et al. (2021) also reported that genetic correlations of microbiota traits with ADG were low compared to RFI, with no estimate significantly different from zero. In addition, an unfavorable genetic correlation between ADG and the Shannon index has also been reported by Lu et al. (2018). Altogether, it suggests that microbiota genetically linked with the host and favoring efficient use of the nutrients, at the digestive and metabolic levels, would slightly reduce growth rate. This is counter-intuitive when considering the genetic correlations between feed efficiency traits and ADG, that are usually negative, and the functional bases of these relationships between gut microbiota, efficiency and growth rate deserve further studies. If confirmed, this antagonistic relationship should deserve further studies to control undesired responses on growth if microbiota was used to improve efficiency.

#### Towards functional hypotheses

The discussion of the role of specific genera and the interpretation of genetic correlations with production traits is generally difficult. For multiple genera, we can find a combination of consistencies and discrepancies between known functions of some microbiota, or previously reported associations with feed efficiency, and the direction of the correlations observed in our dataset. In addition, some of those genera were found associated with efficiency in pigs in previous studies applied to pigs of various ages and feeding conditions, with in some cases opposite associations or no indication of the direction, as already reported in the review of Gardiner et al. (2020). For instance, some of the top associated genera of the two groups are known to produce short chain fatty acids (SCFA) from fiber digestion, which was initially claimed to favor efficiency. For instance, in the *Lachnospiraceae* family *Ruminococcaceae_UCG0002* and *Oscillospira* (Gophna et al. 2017) were associated to better efficiency in our analyses, but *Butyricicoccus* (Quan et al., 2020) *Prevotella_2* (Shah and Collins, 1990) and *Alloprevotella* (Downes et al. 2013) were associated with decreased efficiency. Two hypotheses can be considered: 1) the microbiota functions involved in fibers degradations are not expressed similarly in the genera of the different groups, or 2) (some of) these genera compete with their host for the use of some nutrients, degrading their efficiency, as proposed by Gardiner et al. (2020). At the genetic level, this competition could be enhanced in pigs having genetic abilities to better absorb and metabolize nutrients.

Alternatively, the microbiota diversity and composition may reflect differences in terms of genetically driven host features impacting digestive efficiency. It has been recently shown in humans and humanized mice that the gut microbiota richness and enterotypes were influenced by the intestinal transit time, especially the colon transit time (Kashyap et al. 2013; Vandeputte et al. 2016). Interestingly, Vandeputte et al. (2016) also found, in humans, two main enterotypes dominated either by the *Prevotella* or the *Ruminococcaceae*-*Bacteroides* (RB) genera. Women belonging to the *Prevotella* enterotype had on average looser stools than the ones from the RB enterotype, indicating a faster transit time. These authors hypothesized that gut transit time could be a biological mechanism shaping the gut microbiota through selective pressure on microbial life-strategies, privileging either bacteria with faster growth rates or other properties, as for instance adherence to host tissues, to avoid washout when transit time is faster. As suggested by Déru et al. (2021a), the antagonism between ADG and digestive efficiency may be due to the faster rate of passage in the gut intestinal tract because animals with higher growth rate also have increased DFI. Following this rationale, gut microbiota composition could be a proxy of intestinal transit time, a major determinant of digestive efficiency.

#### Potential for pig selection on microbiota information

To improve feed and digestive efficiency traits that are costly to measure, the present study shows that an alternative to direct selection could be to select genera abundances or an alpha diversity index, which have sufficient heritability and high genetic correlations with efficiency traits. With the two tested diets, we showed limited genetic x diets interactions on the microbiota traits, which suggests that the same selection effort can be used to address the needs of different production systems, for instance. These results should however be consolidated with a wider range of diets. Among the alpha diversity index, the Shannon index seems particularly interesting, as it has the highest heritability among the four indices and is favorably correlated with digestive and feed efficiency traits. In addition, selecting animals on microbiota diversity could have many other positive impacts in relation to improved gut health, such as preventing post-weaning diarrhea and reducing the use of antibiotics (Fouhse et al., 2016). Compared to the selection of one or few microbiota, it would limit risks of specializing animals for certain diets or conditions, or the risk of unbalancing gut microbiota populations.

However, analysis of microbiota samples on farm is more complex than a NIRS-based digestibility predictions because microbiota samples must be frozen immediately in liquid nitrogen and then stored at -80°C, while storing the sample at – 20°C in the freezer is sufficient for the NIRS method. However, the cost was very close for microbiota analysis and a NIRS-based digestibility predictions. Eighty percent of the cost for determining microbiota composition from fecal samples (excluding labor required for fecal collection and material collection) is related to sequencing depth and 20% to microbial DNA extraction. A relatively high sequencing depth (here a minimum of 10,000 sequences per sample) is necessary to accurately quantify genera abundances: in our dataset, 79% of the genera had less than 100 counts on average, 90% had less than 200 counts, which corresponds to only 6 (CO) and 7 (HF) genera with more than 200 counts on average. The 10,000 reads thus corresponds to a minimum cost of phenotyping for the selection of some genera strongly correlated with digestive efficiency traits, as most are not among the most frequent ones. However, it is possible that variance components of the Shannon index would remain stable with lower sequencing depth, thus reducing sequencing cost, and this option could be considered in the future.

In conclusion, gut microbiota analysis is a promising approach to improve the feed and digestive efficiency of growing pigs, which could be applied to pigs fed a range of diets from CO to alternative diets including more dietary fibers. Interactions between host genotype and diets seem limited on the gut microbiota composition. Some microbiota traits, including two diversity indices, could be used as proxy for digestive or feed efficiency in genetic selection, irrespective of the diet. Groups of genera were genetically correlated to each other, reflecting compositional nature of the microbiota data. Thus, it is likely that selection on targeted genera would have an impact on other genera but also on metabolic pathways, and additional work would be needed to investigate these dynamics.

## List of abbreviations

ADG: average daily gain
CO: conventional
DC: digestibility coefficient
DFI: daily feed intake
FCR: feed conversion ratio
HF: high fiber
LW: Large White
Microbiota traits: microbiota genera + alpha diversity indices
NDF: neutral detergent fiber
NE: net energy
NIRS: near infrared spectrometry
OTU: operational taxonomic unit
PCR: polymerase chain reaction
RFI: residual feed intake

