## Supplementary Table S1 for "Genetic relationships between efficiency traits and gut microbiota traits in growing pigs fed a conventional or a high fiber diet"

**Supplementary Table S1**: Ingredient composition (%) of the conventional (CO) and the high fiber (HF) diets

| | Growing phase | | Finishing phase | |
|---|---|---|---|---|
| Ingredients (%) | CO diet | HF diet | CO diet | HF diet |
| Wheat | 38.29 | 38.00 | 42.57 | 39.30 |
| Corn | 25.00 | 0.00 | 25.00 | 0.00 |
| Barley | 15.00 | 16.87 | 15.00 | 17.60 |
| Rapeseed meal | 6.00 | 6.00 | 10.00 | 9.90 |
| Sunflower meal no shelled | 3.00 | 3.00 | 4.80 | 3.00 |
| Soybean meal. 48% CP | 10.4 | 5.40 | 2.50 | 0.00 |
| Wheat bran | 0.00 | 15.00 | 0.00 | 15.00 |
| Soybean hulls | 0.00 | 8.00 | 0.00 | 8.00 |
| Beet pulp | 0.00 | 5.00 | 0.00 | 5.00 |
| L-Lys HCL | 0.44 | 0.35 | 0.11 | 0.31 |
| DL-Met | 0.09 | 0.03 | 0.01 | 0.00 |
| L-Thr | 0.13 | 0.11 | 0.02 | 0.10 |
| Pure valine | 0.02 | 0.00 | 0.00 | 0.00 |
| Calcium carbonate | 1.40 | 1.12 | 0.12 | 1.01 |
| Dicalcium phosphate | 0.49 | 0.29 | 0.05 | 0.00 |
| NaCl | 0.40 | 0.40 | 0.40 | 0.40 |
| Vitamin and trace mineral mixture | 0.40 | 0.40 | 0.40 | 0.40 |
