## Supplementary Table S2 for "Genetic relationships between efficiency traits and gut microbiota traits in growing pigs fed a conventional or a high fiber diet"

**Supplementary Table S2**. Relative abundances of the genera (log10‱) in the population and lines

| | Total | | Conventional diet | | High fiber diet | | |
|---|---|---|---|---|---|---|---|
| | Mean | SD | Mean | SD | Mean | SD | $P^1$ diet |
| *Agathobacter* | 2.41 | 0.28 | 2.33 | 0.28 | 2.50 | 0.24 | <0.001 |
| *Alloprevotella* | 2.09 | 0.24 | 2.05 | 0.26 | 2.14 | 0.20 | <0.001 |
| *Anaerostipes* | 0.87 | 0.52 | 0.74 | 0.50 | 1.01 | 0.50 | <0.001 |
| *Anaerovibrio* | 1.27 | 0.62 | 1.07 | 0.64 | 1.48 | 0.53 | <0.001 |
| *Blautia* | 2.40 | 0.26 | 2.43 | 0.22 | 2.37 | 0.29 | <0.001 |
| *Butyricicoccus* | 1.14 | 0.37 | 1.16 | 0.37 | 1.11 | 0.36 | <0.001 |
| *Campylobacter* | 1.24 | 0.46 | 1.21 | 0.45 | 1.27 | 0.47 | 0.004 |
| *Candidatus_Soleaferrea* | 0.69 | 0.43 | 0.81 | 0.39 | 0.56 | 0.43 | <0.001 |
| *Christensenellaceae_R7_group* | 1.42 | 0.45 | 1.54 | 0.43 | 1.29 | 0.44 | <0.001 |
| *Clostridium_sensu_stricto_1* | 2.45 | 0.41 | 2.61 | 0.38 | 2.28 | 0.36 | <0.001 |
| *Coprococcus_1* | 1.39 | 0.46 | 1.34 | 0.49 | 1.44 | 0.42 | <0.001 |
| *Coprococcus_2* | 1.35 | 0.45 | 1.13 | 0.42 | 1.60 | 0.34 | <0.001 |
| *Coprococcus_3* | 1.94 | 0.26 | 1.97 | 0.25 | 1.92 | 0.26 | <0.001 |
| *Desulfovibrio* | 0.87 | 0.40 | 0.94 | 0.36 | 0.79 | 0.42 | <0.001 |
| *Dialister* | 1.88 | 0.43 | 1.96 | 0.40 | 1.78 | 0.44 | <0.001 |
| *Dorea* | 1.73 | 0.27 | 1.78 | 0.25 | 1.68 | 0.29 | <0.001 |
| *Faecalibacterium* | 2.17 | 0.34 | 2.12 | 0.33 | 2.22 | 0.35 | <0.001 |
| *Family_XIII_AD3011_group* | 1.30 | 0.35 | 1.43 | 0.29 | 1.15 | 0.34 | <0.001 |
| *Family_XIII_UCG001* | 1.00 | 0.26 | 1.03 | 0.27 | 0.98 | 0.25 | <0.001 |
| *Fournierella* | 0.99 | 0.37 | 0.99 | 0.38 | 0.99 | 0.35 | 0.67 |
| *Fusicatenibacter* | 1.34 | 0.35 | 1.39 | 0.33 | 1.29 | 0.37 | <0.001 |
| *Intestinibacter* | 1.59 | 0.21 | 1.60 | 0.22 | 1.58 | 0.21 | 0.046 |
| *Intestinimonas* | 0.69 | 0.39 | 0.77 | 0.35 | 0.60 | 0.41 | <0.001 |
| *Lachnoclostridium* | 1.01 | 0.37 | 0.96 | 0.39 | 1.06 | 0.35 | <0.001 |
| *Lachnospira* | 1.85 | 0.37 | 1.61 | 0.31 | 2.10 | 0.24 | <0.001 |
| *Lachnospiraceae_AC2044_group* | 1.01 | 0.53 | 0.94 | 0.52 | 1.09 | 0.53 | <0.001 |
| *Lachnospiraceae_FCS020_group* | 1.11 | 0.28 | 1.09 | 0.28 | 1.13 | 0.28 | 0.001 |
| *Lachnospiraceae_NC2004_group* | 0.86 | 0.47 | 0.63 | 0.42 | 1.10 | 0.40 | <0.001 |
| *Lachnospiraceae_ND3007_group* | 1.39 | 0.32 | 1.30 | 0.34 | 1.48 | 0.28 | <0.001 |
| *Lachnospiraceae_NK3A20_group* | 0.84 | 0.44 | 0.89 | 0.41 | 0.77 | 0.45 | <0.001 |
| *Lachnospiraceae_NK4A136_group* | 1.63 | 0.33 | 1.53 | 0.29 | 1.74 | 0.32 | <0.001 |
| *Lachnospiraceae_NK4B4_group* | 0.68 | 0.40 | 0.77 | 0.40 | 0.59 | 0.37 | <0.001 |
| *Lachnospiraceae_UCG001* | 1.30 | 0.44 | 1.12 | 0.45 | 1.50 | 0.33 | <0.001 |
| *Lachnospiraceae_UCG004* | 0.61 | 0.38 | 0.59 | 0.39 | 0.63 | 0.36 | 0.09 |
| *Lactobacillus* | 3.03 | 0.38 | 3.02 | 0.40 | 3.05 | 0.35 | 0.14 |
| *Marvinbryantia* | 1.59 | 0.33 | 1.46 | 0.30 | 1.73 | 0.30 | <0.001 |

| | | | | | | | |
|---|---|---|---|---|---|---|---|
| *Mitsuokella* | 1.35 | 0.42 | 1.41 | 0.47 | 1.30 | 0.35 | <0.001 |
| *Mogibacterium* | 1.00 | 0.32 | 1.09 | 0.30 | 0.90 | 0.32 | <0.001 |
| *Oribacterium* | 1.40 | 0.32 | 1.46 | 0.30 | 1.35 | 0.32 | <0.001 |
| *Oscillibacter* | 0.91 | 0.40 | 0.85 | 0.39 | 0.97 | 0.38 | <0.001 |
| *Oscillospira* | 1.11 | 0.34 | 1.05 | 0.36 | 1.17 | 0.30 | <0.001 |
| *Parabacteroides* | 0.99 | 0.47 | 1.09 | 0.43 | 0.89 | 0.49 | <0.001 |
| *Peptococcus* | 0.88 | 0.34 | 0.95 | 0.32 | 0.79 | 0.35 | <0.001 |
| *Prevotella_1* | 1.84 | 0.46 | 1.73 | 0.46 | 1.97 | 0.43 | <0.001 |
| *Prevotella_2* | 1.82 | 0.29 | 1.88 | 0.30 | 1.77 | 0.7 | <0.001 |
| *Prevotella_7* | 2.02 | 0.43 | 2.07 | 0.41 | 1.96 | 0.45 | <0.001 |
| *Prevotella_9* | 3.01 | 0.25 | 3.00 | 0.26 | 3.02 | 0.24 | 0.09 |
| *Prevotellaceae_NK3B31_group* | 2.32 | 0.46 | 2.17 | 0.45 | 2.48 | 0.42 | <0.001 |
| *Prevotellaceae_UCG003* | 1.31 | 0.40 | 1.35 | 0.38 | 1.26 | 0.42 | <0.001 |
| *Rikenellaceae_RC9_gut_group* | 1.96 | 0.28 | 2.01 | 0.27 | 1.91 | 0.27 | <0.001 |
| *Romboutsia* | 1.01 | 0.51 | 1.07 | 0.49 | 0.94 | 0.51 | <0.001 |
| *Roseburia* | 1.87 | 0.38 | 1.71 | 0.36 | 2.04 | 0.31 | <0.001 |
| *Ruminiclostridium_5* | 1.04 | 0.30 | 1.10 | 0.29 | 0.97 | 0.30 | <0.001 |
| *Ruminiclostridium_6* | 1.08 | 0.38 | 1.05 | 0.36 | 1.12 | 0.39 | <0.001 |
| *Ruminiclostridium_9* | 0.98 | 0.27 | 0.99 | 0.27 | 0.96 | 0.28 | 0.01 |
| *Ruminococcaceae_NK4A214_group* | 1.42 | 0.32 | 1.51 | 0.29 | 1.33 | 0.32 | <0.001 |
| *Ruminococcaceae_UCG002* | 1.65 | 0.34 | 1.80 | 0.28 | 1.49 | 0.33 | <0.001 |
| *Ruminococcaceae_UCG005* | 2.08 | 0.33 | 2.05 | 0.31 | 2.12 | 0.35 | <0.001 |
| *Ruminococcaceae_UCG008* | 1.92 | 0.28 | 2.00 | 0.32 | 1.83 | 0.20 | <0.001 |
| *Ruminococcaceae_UCG010* | 1.25 | 0.41 | 1.38 | 0.37 | 1.10 | 0.40 | <0.001 |
| *Ruminococcaceae_UCG013* | 0.80 | 0.37 | 0.76 | 0.36 | 0.90 | 0.36 | <0.001 |
| *Ruminococcaceae_UCG014* | 1.82 | 0.26 | 1.79 | 0.26 | 1.85 | 0.26 | <0.001 |
| *Ruminococcus_1* | 2.24 | 0.24 | 2.12 | 0.20 | 2.38 | 0.20 | <0.001 |
| *Ruminococcus_2* | 1.53 | 0.31 | 1.54 | 0.31 | 1.51 | 0.30 | 0.03 |
| *Shuttleworthia* | 1.66 | 0.65 | 1.77 | 0.64 | 1.54 | 0.65 | <0.001 |
| *Streptococcus* | 2.87 | 0.36 | 2.96 | 0.35 | 2.76 | 0.35 | <0.001 |
| *Subdoligranulum* | 2.14 | 0.21 | 2.15 | 0.20 | 2.12 | 0.22 | 0.048 |
| *Succinivibrio* | 1.53 | 0.50 | 1.65 | 0.47 | 1.40 | 0.51 | <0.001 |
| *Terrisporobacter* | 2.20 | 0.35 | 2.29 | 0.34 | 2.10 | 0.35 | <0.001 |
| *Treponema_2* | 1.50 | 0.49 | 1.48 | 0.48 | 1.51 | 0.51 | 0.18 |
| *Turicibacter* | 0.94 | 0.51 | 1.04 | 0.50 | 0.82 | 0.50 | <0.001 |

[1] P-value obtained from the Kolmogorov-Smirnov test
