## Supplementary Table S3 for "Genetic relationships between efficiency traits and gut microbiota traits in growing pigs fed a conventional or a high fiber diet"

**Supplementary Table S3**. Heritability and confidence intervals for seventy-one genera (at the top of the table. in white) and four alpha diversity indices (at the bottom of the table. in grey) for data of growing pigs fed a conventional (CO) diet and a high fiber (HF) diet analysed jointly. and then separately. and genetic correlations (standard errors) between diets for genera with heritability significantly different from zero in the two diets.

| | Both diets | | CO diet | | | HF diet | | | Genetic correlation[1] |
|---|---|---|---|---|---|---|---|---|---|
| | $h^2$ | CI 95% | $h^2$ | CI 95% | CI 90% | $h^2$ | CI 95% | CI 90% | |
| *Agathobacter* | 0.10 | [0.00-0.20] | *0.00* | [0.00-0.13] | [0.00-0.09] | 0.18 | [0.00-0.38] | [0.02-0.34] | |
| *Alloprevotella* | 0.08 | [0.00-0.18] | 0.13 | [0.00-0.33] | [0.00-0.29] | 0.14 | [0.00-0.34] | [0.00-0.30] | 0.47 (0.43) |
| *Anaerostipes* | 0.15 | [0.03-0.27] | 0.14 | [0.00-0.34] | [0.00-0.30] | 0.19 | [0.00-0.41] | [0.01-0.37] | 0.99 (NE) |
| *Anaerovibrio* | 0.31 | [0.17-0.45] | 0.34 | [0.12-0.56] | [0.16-0.52] | 0.35 | [0.10-0.60] | [0.14-0.56] | 0.82 (0.19) |
| *Blautia* | 0.29 | [0.17-0.41] | 0.24 | [0.04-0.44] | [0.08-0.40] | *0.12* | [0.00-0.14] | [0.01-0.24] | |
| *Butyricicoccus* | 0.10 | [0.00-0.20] | *0.11* | [0.00-0.13] | [0.01-0.21] | *0.09* | [0.00-0.14] | [0.00-0.10] | |
| *Campylobacter* | 0.19 | [0.07-0.31] | 0.35 | [0.13-0.57] | [0.17-0.53] | *0.01* | [0.00-0.14] | [0.00-0.10] | |
| *Candidatus_Soleaferrea* | 0.18 | [0.06-0.30] | 0.17 | [0.00-0.37] | [0.01-0.33] | 0.25 | [0.01-0.49] | [0.05-0.45] | 0.75 (0.32) |
| *Christensenellaceae_R7_group* | 0.27 | [0.15-0.39] | 0.29 | [0.07-0.51] | [0.11-0.47] | 0.18 | [0.00-0.40] | [0.00-0.36] | 0.99 (NE) |
| *Clostridium_sensu_stricto_1* | 0.23 | [0.13-0.33] | 0.18 | [0.00-0.36] | [0.03-0.33] | 0.39 | [0.15-0.63] | [0.19-0.59] | 0.99 (NE) |
| *Coprococcus_1* | 0.23 | [0.11-0.35] | 0.20 | [0.00-0.40] | [0.04-0.36] | *0.07* | [0.00-0.14] | [0.00-0.10] | |
| *Coprococcus_2* | 0.10 | [0.00-0.20] | 0.13 | [0.00-0.33] | [0.00-0.29] | *0.00* | [0.00-0.14] | [0.00-0.10] | |
| *Coprococcus_3* | 0.08 | [0.00-0.18] | 0.13 | [0.00-0.31] | [0.00-0.28] | 0.28 | [0.03-0.53] | [0.07-0.49] | 0.51 (0.34) |
| *Desulfovibrio* | 0.19 | [0.07-0.31] | 0.15 | [0.00-0.35] | [0.00-0.31] | *0.08* | [0.00-0.14] | [0.00-0.10] | |
| *Dialister* | 0.30 | [0.18-0.42] | *0.06* | [0.00-0.13] | [0.00-0.09] | 0.33 | [0.11-0.55] | [0.15-0.51] | |
| *Dorea* | 0.15 | [0.03-0.27] | *0.08* | [0.00-0.13] | [0.00-0.09] | 0.17 | [0.00-0.39] | [0.00-0.35] | |

| | | | | | | | | | |
|---|---|---|---|---|---|---|---|---|---|
| *Faecalibacterium* | 0.11 | [0.01-0.21] | *0.02* | [0.00-0.13] | [0.00-0.09] | *0.11* | [0.00-0.14] | [0.00-0.23] | |
| *Family_XIII_AD3011_group* | 0.18 | [0.06-0.30] | 0.21 | [0.03-0.39] | [0.06-0.36] | 0.25 | [0.01-0.49] | [0.05-0.45] | 0.96 (0.34) |
| *Family_XIII_UCG001* | 0.16 | [0.04-0.28] | 0.15 | [0.00-0.35] | [0.00-0.31] | *0.03* | [0.00-0.14] | [0.00-0.10] | |
| *Fournierella* | 0.08 | [0.00-0.18] | *0.11* | [0.00-0.13] | [0.01-0.21] | *0.00* | [0.00-0.14] | [0.00-0.10] | |
| *Fusicatenibacter* | 0.22 | [0.10-0.34] | *0.04* | [0.00-0.13] | [0.00-0.09] | 0.35 | [0.11-0.59] | [0.15-0.55] | |
| *Intestinibacter* | *0.04* | [0.00-0.06] | *0.00* | [0.00-0.13] | [0.00-0.09] | *0.02* | [0.00-0.14] | [0.00-0.10] | |
| *Intestinimonas* | 0.10 | [0.00-0.20] | *0.03* | [0.00-0.13] | [0.00-0.09] | 0.19 | [0.00-0.41] | [0.01-0.37] | |
| *Lachnoclostridium* | 0.05 | [0.00-0.06] | *0.01* | [0.00-0.13] | [0.00-0.09] | *0.00* | [0.00-0.14] | [0.00-0.10] | |
| *Lachnospira* | 0.00 | [0.00-0.06] | *0.00* | [0.00-0.13] | [0.00-0.09] | *0.02* | [0.00-0.14] | [0.00-0.10] | |
| *Lachnospiraceae_AC2044_group* | 0.13 | [0.03-0.23] | 0.14 | [0.00-0.32] | [0.00-0.29] | *0.03* | [0.00-0.14] | [0.00-0.10] | |
| *Lachnospiraceae_FCS020_group* | *0.00* | [0.00-0.06] | *0.00* | [0.00-0.13] | [0.00-0.09] | *0.00* | [0.00-0.14] | [0.00-0.10] | |
| *Lachnospiraceae_NC2004_group* | *0.00* | [0.00-0.06] | *0.00* | [0.00-0.13] | [0.00-0.09] | *0.04* | [0.00-0.14] | [0.00-0.10] | |
| *Lachnospiraceae_ND3007_group* | *0.06* | [0.00-0.14] | *0.12* | [0.00-0.13] | [0.02-0.22] | *0.00* | [0.00-0.14] | [0.00-0.10] | |
| *Lachnospiraceae_NK3A20_group* | 0.20 | [0.08-0.32] | 0.30 | [0.08-0.52] | [0.12-0.48] | 0.29 | [0.07-0.51] | [0.11-0.47] | 0.46 (0.23)[2] |
| *Lachnospiraceae_NK4A136_group* | 0.08 | [0.00-0.18] | *0.08* | [0.00-0.13] | [0.00-0.09] | *0.04* | [0.00-0.14] | [0.00-0.10] | |
| *Lachnospiraceae_NK4B4_group* | 0.08 | [0.00-0.16] | 0.14 | [0.00-0.32] | [0.00-0.29] | *0.02* | [0.00-0.14] | [0.00-0.10] | |
| *Lachnospiraceae_UCG001* | *0.06* | [0.00-0.12] | *0.00* | [0.00-0.13] | [0.00-0.09] | *0.02* | [0.00-0.14] | [0.00-0.10] | |
| *Lachnospiraceae_UCG004* | *0.03* | [0.00-0.06] | *0.01* | [0.00-0.13] | [0.00-0.09] | *0.00* | [0.00-0.14] | [0.00-0.10] | |
| *Lactobacillus* | 0.18 | [0.06-0.30] | *0.10* | [0.00-0.13] | [0.00-0.22] | 0.19 | [0.00-0.43] | [0.00-0.39] | |
| *Marvinbryantia* | 0.06 | [0.04-0.08] | *0.09* | [0.00-0.13] | [0.00-0.09] | *0.00* | [0.00-0.14] | [0.00-0.10] | |
| *Mitsuokella* | 0.18 | [0.06-0.30] | *0.11* | [0.00-0.13] | [0.01-0.21] | *0.05* | [0.00-0.14] | [0.00-0.10] | |
| *Mogibacterium* | 0.14 | [0.04-0.24] | 0.23 | [0.03-0.43] | [0.07-0.39] | 0.15 | [0.00-0.35] | [0.00-0.31] | 0.36 (0.34)[2] |
| *Oribacterium* | 0.19 | [0.07-0.31] | *0.12* | [0.00-0.13] | [0.02-0.22] | 0.20 | [0.00-0.40] | [0.04-0.36] | |
| *Oscillibacter* | 0.15 | [0.03-0.27] | *0.08* | [0.00-0.13] | [0.00-0.09] | *0.13* | [0.00-0.14] | [0.02-0.25] | |
| *Oscillospira* | 0.07 | [0.00-0.15] | *0.12* | [0.00-0.13] | [0.00-0.27] | *0.13* | [0.00-0.14] | [0.02-0.25] | |
| *Parabacteroides* | 0.17 | [0.05-0.29] | 0.17 | [0.00-0.37] | [0.01-0.33] | 0.14 | [0.00-0.34] | [0.00-0.30] | 0.99 (NE) |
| *Peptococcus* | 0.08 | [0.00-0.18] | *0.00* | [0.00-0.13] | [0.00-0.09] | 0.23 | [0.03-0.43] | [0.07-0.39] | |
| *Prevotella_1* | 0.15 | [0.03-0.27] | 0.22 | [0.00-0.44] | [0.04-0.40] | *0.10* | [0.00-0.14] | [0.00-0.22] | |

| | | | | | | | | | |
|---|---|---|---|---|---|---|---|---|---|
| *Prevotella_2* | 0.08 | [0.00-0.18] | *0.00* | [0.00-0.13] | [0.00-0.09] | *0.08* | [0.00-0.14] | [0.00-0.10] | |
| *Prevotella_7* | 0.23 | [0.11-0.35] | 0.22 | [0.02-0.42] | [0.06-0.38] | 0.18 | [0.00-0.38] | [0.02-0.34] | 0.99 (NE) |
| *Prevotella_9* | 0.12 | [0.00-0.24] | *0.00* | [0.00-0.13] | [0.00-0.09] | *0.11* | [0.00-0.14] | [0.00-0.23] | |
| *Prevotellaceae_NK3B31_group* | 0.24 | [0.10-0.38] | 0.17 | [0.00-0.37] | [0.01-0.33] | 0.21 | [0.01-0.41] | [0.05-0.37] | 0.92 (0.21) |
| *Prevotellaceae_UCG003* | 0.26 | [0.12-0.40] | *0.11* | [0.00-0.13] | [0.01-0.21] | 0.31 | [0.07-0.55] | [0.11-0.51] | |
| *Rikenellaceae_RC9_gut_group* | 0.30 | [0.16-0.44] | 0.24 | [0.04-0.44] | [0.08-0.40] | 0.21 | [0.00-0.43] | [0.03-0.39] | 0.99 (NE) |
| *Romboutsia* | 0.26 | [0.14-0.38] | 0.29 | [0.09-0.49] | [0.13-0.45] | 0.30 | [0.08-0.52] | [0.12-0.48] | 0.80 (0.21) |
| *Roseburia* | *0.02* | [0.00-0.06] | *0.04* | [0.00-0.13] | [0.00-0.09] | *0.03* | [0.00-0.14] | [0.00-0.10] | |
| *Ruminiclostridium_5* | 0.10 | [0.00-0.20] | *0.12* | [0.00-0.13] | [0.00-0.25] | 0.14 | [0.00-0.34] | [0.00-0.30] | |
| *Ruminiclostridium_6* | 0.13 | [0.03-0.23] | 0.13 | [0.00-0.31] | [0.00-0.28] | 0.25 | [0.03-0.47] | [0.07-0.43] | 0.58 (0.38) |
| *Ruminiclostridium_9* | *0.00* | [0.00-0.06] | *0.04* | [0.00-0.13] | [0.00-0.09] | *0.00* | [0.00-0.14] | [0.00-0.10] | |
| *Ruminococcaceae_NK4A214_group* | 0.26 | [0.12-0.40] | 0.20 | [0.00-0.42] | [0.02-0.38] | 0.35 | [0.10-0.6] | [0.14-0.56] | 0.99 (NE) |
| *Ruminococcaceae_UCG002* | 0.21 | [0.09-0.33] | *0.06* | [0.00-0.13] | [0.00-0.09] | 0.31 | [0.07-0.55] | [0.11-0.51] | |
| *Ruminococcaceae_UCG005* | 0.23 | [0.11-0.35] | *0.12* | [0.00-0.13] | [0.00-0.27] | 0.18 | [0.00-0.40] | [0.00-0.36] | |
| *Ruminococcaceae_UCG008* | 0.08 | [0.00-0.18] | 0.13 | [0.00-0.31] | [0.00-0.28] | *0.00* | [0.00-0.14] | [0.00-0.10] | |
| *Ruminococcaceae_UCG010* | 0.28 | [0.16-0.40] | 0.25 | [0.05-0.45] | [0.09-0.41] | 0.24 | [0.00-0.48] | [0.04-0.44] | 0.99 (NE) |
| *Ruminococcaceae_UCG013* | *0.05* | [0.00-0.06] | *0.02* | [0.00-0.13] | [0.00-0.09] | *0.04* | [0.00-0.14] | [0.00-0.10] | |
| *Ruminococcaceae_UCG014* | 0.10 | [0.00-0.20] | *0.11* | [0.00-0.13] | [0.01-0.21] | *0.10* | [0.00-0.14] | [0.00-0.22] | |
| *Ruminococcus_1* | 0.14 | [0.04-0.24] | 0.16 | [0.00-0.34] | [0.01-0.31] | 0.17 | [0.00-0.39] | [0.00-0.35] | 0.96 (0.52) |
| *Ruminococcus_2* | 0.08 | [0.00-0.16] | 0.14 | [0.00-0.32] | [0.00-0.29] | 0.23 | [0.01-0.45] | [0.05-0.41] | 0.14 (0.37) [2] |
| *Shuttleworthia* | 0.13 | [0.03-0.23] | 0.16 | [0.00-0.36] | [0.00-0.32] | *0.12* | [0.00-0.14] | [0.01-0.24] | |
| *Streptococcus* | 0.22 | [0.100.34] | 0.19 | [0.01-0.37] | [0.04-0.34] | 0.26 | [0.06-0.46] | [0.10-0.42] | 0.77 (0.22) |
| *Subdoligranulum* | *0.00* | [0.00-0.06] | *0.00* | [0.00-0.13] | [0.00-0.09] | *0.02* | [0.00-0.14] | [0.00-0.10] | |
| *Succinivibrio* | 0.09 | [0.00-0.19] | *0.04* | [0.00-0.13] | [0.00-0.09] | *0.02* | [0.00-0.14] | [0.00-0.10] | |
| *Terrisporobacter* | 0.27 | [0.15-0.39] | 0.18 | [0.00-0.36] | [0.03-0.33] | 0.35 | [0.11-0.59] | [0.15-0.55] | 0.99 (NE) |
| *Treponema_2* | 0.22 | [0.10-0.34] | 0.38 | [0.14-0.62] | [0.18-0.58] | 0.24 | [0.02-0.46] | [0.06-0.42] | 0.70 (0.25) |
| *Turicibacter* | 0.18 | [0.08-0.28] | 0.18 | [0.00-0.38] | [0.02-0.34] | 0.22 | [0.02-0.42] | [0.06-0.38] | 0.80 (0.30) |
| Chao1 | *0.05* | [0.00-0.06] | *0.15* | [0.00-0.33] | [0.00-0.30] | *0.00* | [0.00-0.14] | [0.00-0.10] | |

| | | | | | | | | | |
|---|---|---|---|---|---|---|---|---|---|
| Richness | *0.09* | [0.00-0.19] | *0.15* | [0.00-0.33] | [0.00-0.30] | *0.00* | [0.00-0.14] | [0.00-0.10] | |
| Shannon | 0.19 | [0.09-0.29] | 0.23 | [0.03-0.43] | [0.07-0.39] | 0.20 | [0.06-0.34] | [0.08-0.32] | 0.83 (0.28) |
| Simpson | 0.13 | [0.03-0.23] | *0.19* | [0.00-0.39] | [0.03-0.35] | *0.18* | [0.00-0.38] | [0.02-0.34] | 0.62 (0.32) |

Heritability estimates in italic did not differ from 0.0 (P > 0.05)

[1] NE = non estimable

[2] $P < 0.05$ for the likelihood ratio test to reject the null hypothesis “the genetic correlation is 0.99”
