## Supplementary Table S4 for "Genetic relationships between efficiency traits and gut microbiota traits in growing pigs fed a conventional or a high fiber diet"

**Supplementary Table S4**. Genetic correlations ($r_g$) and their standard errors (se[1]) between genera with heritability significantly different from zero in analyses of both diets together and production traits.

| | ADG | | DFI | | RFI | | FCR | | DC E | | DC OM | | DC N | |
|---|---|---|---|---|---|---|---|---|---|---|---|---|---|---|
| | $r_g$ | se | $r_g$ | se | $r_g$ | se | $r_g$ | se | $r_g$ | se | $r_g$ | se | $r_g$ | se |
| *Agathobacter* | 0.24 | 0.21 | 0.27 | 0.20 | 0.45 | 0.21 | 0.24 | 0.22 | -0.53 | 0.22 | -0.55 | 0.21 | -0.47 | 0.22 |
| *Alloprevotella* | 0.32 | 0.22 | 0.60 | 0.21 | 0.68 | 0.24 | 0.66 | 0.25 | -1.00 | 0.00 | -1.00 | 0.00 | -1.00 | 0.00 |
| *Anaerostipes* | 0.32 | 0.19 | 0.33 | 0.18 | -0.02 | 0.20 | 0.25 | 0.19 | -0.50 | 0.23 | -0.46 | 0.23 | -0.58 | 0.22 |
| *Anaerovibrio* | 0.31 | 0.14 | 0.30 | 0.13 | 0.23 | 0.14 | 0.19 | 0.14 | -0.69 | 0.16 | -0.62 | 0.17 | -0.70 | 0.15 |
| *Blautia* | 0.23 | 0.14 | 0.23 | 0.13 | 0.24 | 0.14 | 0.21 | 0.15 | -0.13 | 0.19 | -0.12 | 0.18 | -0.15 | 0.18 |
| *Butyricicoccus* | 0.40 | 0.21 | 0.53 | 0.20 | 0.69 | 0.20 | 0.43 | 0.21 | -0.63 | 0.25 | -0.62 | 0.24 | -0.56 | 0.25 |
| *Campylobacter* | -0.14 | 0.17 | -0.44 | 0.15 | -0.54 | 0.16 | -0.48 | 0.16 | -0.03 | 0.21 | 0.01 | 0.21 | -0.26 | 0.20 |
| *Candidatus_Soleaferrea* | -0.57 | 0.15 | -0.88 | 0.11 | -0.73 | 0.13 | -0.67 | 0.14 | 0.88 | 0.13 | 0.89 | 0.13 | 0.87 | 0.13 |
| *Christensenellaceae_R7_group* | -0.56 | 0.13 | -0.72 | 0.12 | -0.77 | 0.12 | -0.48 | 0.14 | 0.76 | 0.12 | 0.76 | 0.12 | 0.75 | 0.12 |
| *Clostridium_sensu_stricto_1* | -0.27 | 0.15 | -0.40 | 0.13 | -0.22 | 0.14 | -0.33 | 0.14 | 0.60 | 0.15 | 0.58 | 0.15 | 0.52 | 0.14 |
| *Coprococcus_1* | 0.17 | 0.16 | 0.10 | 0.15 | 0.08 | 0.16 | 0.09 | 0.16 | -0.34 | 0.20 | -0.27 | 0.19 | -0.42 | 0.18 |
| *Coprococcus_2* | 0.31 | 0.23 | 0.62 | 0.21 | 0.67 | 0.22 | 0.66 | 0.23 | -0.28 | 0.27 | -0.25 | 0.27 | -0.14 | 0.27 |
| *Coprococcus_3* | -0.43 | 0.23 | -0.44 | 0.22 | -0.42 | 0.24 | -0.08 | 0.23 | 0.25 | 0.27 | 0.29 | 0.26 | 0.24 | 0.27 |
| *Desulfovibrio* | -0.54 | 0.18 | -0.45 | 0.15 | -0.38 | 0.16 | -0.26 | 0.16 | 0.71 | 0.16 | 0.68 | 0.16 | 0.70 | 0.15 |
| *Dialister* | 0.09 | 0.15 | 0.02 | 0.14 | 0.06 | 0.15 | 0.00 | 0.15 | 0.04 | 0.19 | 0.01 | 0.18 | 0.17 | 0.18 |
| *Dorea* | 0.21 | 0.18 | 0.11 | 0.18 | 0.23 | 0.19 | -0.06 | 0.19 | -0.60 | 0.22 | -0.60 | 0.21 | -0.52 | 0.21 |
| *Faecalibacterium* | 0.49 | 0.19 | 0.69 | 0.17 | 0.89 | 0.20 | 0.67 | 0.20 | -0.67 | 0.22 | -0.67 | 0.21 | -0.59 | 0.22 |
| *Family_XIII_AD3011_group* | -0.53 | 0.16 | -0.76 | 0.13 | -0.87 | 0.13 | -0.53 | 0.15 | 1.00 | 0.00 | 1.00 | 0.00 | 1.00 | 0.10 |
| *Family_XIII_UCG001* | -0.17 | 0.19 | -0.39 | 0.16 | -0.41 | 0.16 | -0.45 | 0.16 | 0.75 | 0.18 | 0.70 | 0.18 | 0.82 | 0.16 |
| *Fournierella* | 0.06 | 0.24 | 0.29 | 0.21 | 0.42 | 0.22 | 0.44 | 0.23 | -0.45 | 0.30 | -0.42 | 0.30 | -0.64 | 0.27 |
| *Fusicatenibacter* | 0.25 | 0.16 | 0.47 | 0.13 | 0.53 | 0.13 | 0.47 | 0.14 | -0.38 | 0.19 | -0.38 | 0.18 | -0.47 | 0.17 |
| *Intestinimonas* | -0.06 | 0.22 | -0.19 | 0.19 | -0.04 | 0.21 | -0.23 | 0.21 | -0.27 | 0.28 | -0.28 | 0.27 | -0.28 | 0.27 |
| *Lachnospiraceae_AC2044_group* | -0.53 | 0.18 | -0.18 | 0.19 | -0.02 | 0.20 | 0.17 | 0.19 | 0.19 | 0.24 | 0.22 | 0.24 | 0.00 | 0.24 |
| *Lachnospiraceae_NK3A20_group* | -0.12 | 0.17 | -0.25 | 0.15 | -0.28 | 0.15 | -0.28 | 0.16 | 0.71 | 0.18 | 0.65 | 0.18 | 0.76 | 0.16 |

| | | | | | | | | | | | | | | |
|---|---|---|---|---|---|---|---|---|---|---|---|---|---|---|
| Lachnospiraceae_NK4A136_group | -0.20 | 0.25 | -0.35 | 0.23 | -0.43 | 0.24 | -0.16 | 0.23 | 0.70 | 0.32 | 0.70 | 0.31 | 0.62 | 0.30 |
| Lachnospiraceae_NK4B4_group | -0.42 | 0.26 | -0.44 | 0.23 | -0.27 | 0.23 | -0.35 | 0.22 | 0.39 | 0.29 | 0.35 | 0.28 | 0.46 | 0.27 |
| Lactobacillus | 0.04 | 0.17 | 0.21 | 0.16 | 0.11 | 0.17 | 0.24 | 0.17 | -0.57 | 0.20 | -0.56 | 0.20 | -0.56 | 0.19 |
| Marvinbryantia | 0.28 | 0.23 | 0.07 | 0.23 | -0.01 | 0.25 | 0.02 | 0.25 | -0.09 | 0.31 | -0.04 | 0.30 | -0.27 | 0.29 |
| Mitsuokella | 0.06 | 0.17 | 0.14 | 0.17 | 0.20 | 0.18 | 0.13 | 0.18 | 0.17 | 0.22 | 0.10 | 0.21 | 0.26 | 0.21 |
| Mogibacterium | -0.30 | 0.19 | -0.52 | 0.17 | -0.67 | 0.17 | -0.62 | 0.17 | 0.76 | 0.21 | 0.75 | 0.20 | 0.80 | 0.21 |
| Oribacterium | 0.06 | 0.17 | -0.05 | 0.16 | 0.15 | 0.17 | -0.05 | 0.17 | 0.19 | 0.21 | 0.15 | 0.21 | 0.27 | 0.20 |
| Oscillibacter | -0.57 | 0.21 | -0.69 | 0.17 | -0.62 | 0.18 | -0.40 | 0.17 | 0.88 | 0.20 | 0.90 | 0.19 | 0.81 | 0.19 |
| Oscillospira | 0.03 | 0.26 | -0.53 | 0.26 | -0.45 | 0.27 | -0.54 | 0.23 | 0.92 | 0.37 | 0.97 | 0.38 | 0.90 | 0.36 |
| Parabacteroides | -0.37 | 0.17 | -0.54 | 0.15 | -0.69 | 0.15 | -0.53 | 0.17 | 0.17 | 0.22 | 0.15 | 0.22 | 0.03 | 0.22 |
| Peptococcus | 0.30 | 0.23 | 0.28 | 0.22 | 0.27 | 0.24 | 0.03 | 0.24 | 0.01 | 0.30 | 0.05 | 0.30 | 0.31 | 0.28 |
| Prevotella_1 | -0.21 | 0.21 | -0.51 | 0.18 | -0.39 | 0.19 | -0.36 | 0.18 | 0.47 | 0.22 | 0.45 | 0.22 | 0.34 | 0.22 |
| Prevotella_2 | 0.39 | 0.27 | 0.73 | 0.29 | 1.00 | 0.00 | 0.49 | 0.28 | -0.89 | 0.36 | -0.92 | 0.35 | -0.98 | 0.32 |
| Prevotella_7 | 0.08 | 0.16 | -0.04 | 0.15 | 0.01 | 0.16 | -0.13 | 0.16 | 0.36 | 0.20 | 0.31 | 0.19 | 0.47 | 0.18 |
| Prevotella_9 | 0.30 | 0.19 | 0.45 | 0.17 | 0.68 | 0.18 | 0.43 | 0.20 | -0.59 | 0.20 | -0.62 | 0.19 | -0.50 | 0.20 |
| Prevotellaceae_NK3B31_group | -0.18 | 0.16 | -0.26 | 0.15 | -0.31 | 0.16 | -0.26 | 0.16 | 0.07 | 0.20 | 0.11 | 0.20 | 0.06 | 0.20 |
| Prevotellaceae_UCG003 | -0.30 | 0.15 | -0.32 | 0.14 | -0.41 | 0.15 | -0.29 | 0.15 | 0.33 | 0.19 | 0.32 | 0.18 | 0.19 | 0.19 |
| Rikenellaceae_RC9_gut_group | -0.47 | 0.13 | -0.67 | 0.10 | -0.64 | 0.12 | -0.46 | 0.13 | 0.79 | 0.11 | 0.78 | 0.11 | 0.72 | 0.12 |
| Romboutsia | -0.20 | 0.14 | -0.45 | 0.12 | -0.25 | 0.14 | -0.40 | 0.13 | 0.58 | 0.15 | 0.55 | 0.15 | 0.52 | 0.14 |
| Ruminiclostridium_5 | -0.20 | 0.22 | -0.40 | 0.20 | -0.39 | 0.21 | -0.40 | 0.20 | 0.33 | 0.25 | 0.31 | 0.25 | 0.32 | 0.24 |
| Ruminiclostridium_6 | -0.43 | 0.20 | -0.31 | 0.18 | -0.20 | 0.19 | -0.07 | 0.20 | 0.16 | 0.24 | 0.14 | 0.23 | 0.16 | 0.23 |
| Ruminococcaceae_NK4A214_group | -0.46 | 0.14 | -0.68 | 0.11 | -0.70 | 0.12 | -0.47 | 0.13 | 0.96 | 0.09 | 0.96 | 0.09 | 0.94 | 0.09 |
| Ruminococcaceae_UCG005 | -0.36 | 0.15 | -0.46 | 0.14 | -0.48 | 0.15 | -0.30 | 0.15 | 0.33 | 0.18 | 0.37 | 0.18 | 0.33 | 0.18 |
| Ruminococcaceae_UCG002 | -0.54 | 0.14 | -0.87 | 0.10 | -0.77 | 0.12 | -0.67 | 0.13 | 0.97 | 0.10 | 0.98 | 0.09 | 0.95 | 0.10 |
| Ruminococcaceae_UCG008 | -0.20 | 0.27 | 0.20 | 0.23 | 0.27 | 0.23 | 0.28 | 0.24 | -0.16 | 0.32 | -0.14 | 0.31 | -0.43 | 0.29 |
| Ruminococcaceae_UCG010 | -0.48 | 0.15 | -0.74 | 0.11 | -0.60 | 0.13 | -0.49 | 0.13 | 0.89 | 0.10 | 0.90 | 0.10 | 0.81 | 0.11 |
| Ruminococcaceae_UCG014 | -0.55 | 0.22 | -0.72 | 0.21 | -0.56 | 0.21 | -0.41 | 0.21 | 0.76 | 0.22 | 0.75 | 0.21 | 0.83 | 0.21 |
| Ruminococcus_1 | -0.62 | 0.16 | -0.62 | 0.15 | -0.45 | 0.17 | -0.22 | 0.18 | 0.69 | 0.17 | 0.70 | 0.17 | 0.70 | 0.17 |
| Ruminococcus_2 | 0.34 | 0.21 | 0.37 | 0.20 | 0.33 | 0.22 | 0.30 | 0.23 | -0.37 | 0.29 | -0.35 | 0.29 | -0.47 | 0.28 |
| Shuttleworthia | -0.22 | 0.19 | -0.21 | 0.17 | -0.10 | 0.19 | -0.15 | 0.19 | 0.78 | 0.22 | 0.71 | 0.22 | 0.72 | 0.21 |

| | | | | | | | | | | | | | | |
|---|---|---|---|---|---|---|---|---|---|---|---|---|---|---|
| *Streptococcus* | 0.06 | 0.15 | 0.00 | 0.14 | 0.00 | 0.16 | -0.16 | 0.15 | 0.44 | 0.18 | 0.45 | 0.17 | 0.58 | 0.16 |
| *Succinivibrio* | 0.08 | 0.22 | -0.15 | 0.20 | 0.12 | 0.23 | -0.05 | 0.22 | -0.27 | 0.27 | -0.26 | 0.26 | -0.17 | 0.27 |
| *Terrisporobacter* | -0.33 | 0.14 | -0.49 | 0.11 | -0.28 | 0.13 | -0.39 | 0.12 | 0.62 | 0.14 | 0.59 | 0.14 | 0.54 | 0.14 |
| *Treponema_2* | -0.51 | 0.14 | -0.48 | 0.14 | -0.47 | 0.15 | -0.22 | 0.16 | 0.51 | 0.16 | 0.53 | 0.15 | 0.38 | 0.17 |
| *Turicibacter* | -0.22 | 0.17 | -0.32 | 0.15 | -0.12 | 0.17 | -0.28 | 0.16 | 0.42 | 0.19 | 0.42 | 0.19 | 0.36 | 0.19 |
| Shannon | -0.29 | 0.17 | -0.56 | 0.13 | -0.51 | 0.14 | -0.46 | 0.14 | 0.91 | 0.13 | 0.90 | 0.13 | 0.88 | 0.12 |
| Simpson | -0.19 | 0.19 | -0.48 | 0.16 | -0.42 | 0.17 | -0.42 | 0.17 | 1.00 | 0.00 | 1.00 | 0.00 | 1.00 | 0.00 |
| Richness | -0.87 | 0.27 | -0.81 | 0.20 | -0.61 | 0.20 | -0.55 | 0.18 | 0.74 | 0.24 | 0.70 | 0.24 | 0.58 | 0.24 |

[1] NE=not estimable
